# Convergent and divergent phenotypic impacts of estradiol on neurodevelopmental disorder risk genes

**DOI:** 10.64898/2026.02.18.706588

**Authors:** April Pruitt, Liang Yang, Seoyeon Lee, Novin Balafkan, Shilin Gao, Carina Seah, Yi Dai, Elizabeth Davidson, Suha Khan, Annabel Sen, Jenny Liu, Isabel Wood, Gang Xu, Xiayuan Huang, Marina Carlson, Dejian Zhao, Priyanka Jamadagni, Grace Dossou, Kayla Retallick-Townsley, Xinyun Li, Xenophon Papademetris, P.J. Michael Deans, Meilin Fernandez Garcia, Huanyao Gao, Tianyun Wang, Zuoheng Wang, Laura Huckins, Ellen J. Hoffman, Kristen Brennand

## Abstract

Sex hormone signaling during neurodevelopment may act as a modulator of risk for neurodevelopmental disorders (NDDs), including autism spectrum disorders (ASDs). By conducting a systematic and multi-modal evaluation of estradiol effects across loss-of-function deletions in 36 functionally diverse, large-effect ASD/NDD genes in human induced pluripotent stem cell-derived neurons and larval zebrafish, we uncover convergent and divergent gene-by-estradiol interactions at the transcriptomic, circuit, and behavioral levels. Whereas estradiol partially ameliorates dysregulated expression signatures across all ASD/NDD gene knockouts examined, it selectively and rapidly dampens network hyperexcitability phenotypes in only a subset of ASD/NDD gene knockouts. Two genes, *ASH1L* and *SCN2A*, are particularly responsive to estradiol treatment, with dramatic rescue of molecular and cellular effects in human knockout neurons, coupled with behavioral phenotypes in larval zebrafish mutants. The seven most estradiol responsive genes show enhanced protein-protein interaction network connectivity, strong co-expression in specific postnatal time periods, and enrichment in upper-layer mature excitatory neurons. Altogether, we describe distinct estradiol rescue in different modalities, elucidating shared transcriptomic effects as well as gene-specific cellular and behavioral rescue in the context of loss of function mutations of select ASD/NDD genes.

## INTRODUCTION

Estradiol plays a critical role in the brain^1–3^, where it regulates neurogenesis^4,5^, intrinsic neuronal excitability^6,7^, synaptic plasticity^8–10^, and behavior^11^ across both excitatory^12,13^ and inhibitory^14,15^ circuits. These neuromodulatory effects occur through two cooperative signaling mechanisms. Classic nuclear receptors (ligand-activated membrane estrogen receptors ERα and ERβ) are located within the cytoplasm and, upon estradiol binding, translocate to the nucleus to function as transcription factors and regulate gene transcription over a period of hours to days (“genomic mechanism”)^16^. Brain-specific ERα targets are uniquely enriched for neurotransmitter receptors, ion channels, neurotrophin receptors, as well as ASD/NDD genes^17^; yet many estradiol effects are so rapid that they cannot result from RNA transcription and subsequent protein synthesis^18^. Fast-acting membrane bound receptors (e.g., G-protein coupled receptor GPER1) activate second messengers, kinases, and ion fluxes (e.g., calcium)^19^ to regulate synaptic function in minutes (“non-genomic mechanism”)^14,20^. Neuronal estradiol responses differ significantly between males and females, reflecting variations in receptor expression and distribution as well as sex-specific patterns of gene expression and brain circuitry^17^. Towards resolving if neuronal estradiol responses are otherwise genotype-dependent, genetics studies queried common genetic variation and discovered loci associated with circulating estrogen levels^21,22^ as well as widespread polymorphisms in estrogen response elements (EREs) across the human genome that modulate genomic response to estradiol in a cell-and tissue-specific manner^23,24^.

The phenotypic spectrum and severity of ASD/NDD is associated with the complex interactions of rare mutations^25–27^, polygenic risk^28^, and sex effects^29,30^. At the same time, there is growing evidence from our group and others that estrogens modulate ASD/NDD gene-associated phenotypes in distinct contexts. In an unbiased screen, we identified estrogenic compounds as suppressors of nighttime hyperactivity in zebrafish *CNTNAP2*^31^ mutants. Our group likewise showed that estradiol ameliorated signaling deficits associated with *NRXN1* loss of function mutations in functionally mature patient-derived human induced pluripotent (hiPSC)-neurons^32^. Furthermore, estradiol rescued increased neural cell proliferation in developing tadpoles lacking the function of *DYRK1A*^33^, proliferation deficits in knockdown hiPSC neural progenitor cells (NPCs) (*DYRK1A, NRXN1, ADNP*^33^), and synaptic phenotypes in transgenic mouse neurons (*DISC1*^34^). However, the extent to which estradiol might broadly function as a modulator of ASD/NDD gene-associated phenotypes across genes and neurodevelopmental contexts is currently unknown.

Here, we seek to systematically evaluate whether and how 17β-estradiol, the primary biologically active form of estrogen, shapes phenotypes associated with functionally diverse ASD/NDD risk genes. Utilizing a large-scale, multi-modal functional screening pipeline, we interrogated how estradiol exposure shapes neurodevelopmental molecular, cellular, and behavioral phenotypic differences across 36 functionally diverse and highly penetrant ASD/NDD loss-of-function (LOF) genetic mutations (*ANK2, ANK3, ARID1B, ASH1L, ASXL3, BCL11A, CACNA1G, CHD2, CHD8, CREBBP, CUL3, DPYSL2, DYRK1A, FOXP2, GRIA3, GRIN2A, KDM5B, KDM6B, KMT2C, KMT5B, MBD5, MED13L, NRXN1, PHF12, PHF21A, POGZ, PPP2R5D, PTEN, SCN2A, SETD5, SHANK3, SIN3A, SKI, SLC6A1, SMARCC2, WAC)* (**Fig. 1**). Mutations in these large-effect ASD/NDD risk genes perturb key early developmental pathways, such as transcriptional regulation, neurogenesis, synaptic maturation, and circuit maturation^35–41^, which are also sensitive to estradiol modulation. Therefore, interrogating the functional readout of the interaction of ASD/NDD gene mutations and estradiol may provide greater mechanistic insight into regulation of essential neuronal function and developmental phenotypes.

**Figure 1:**
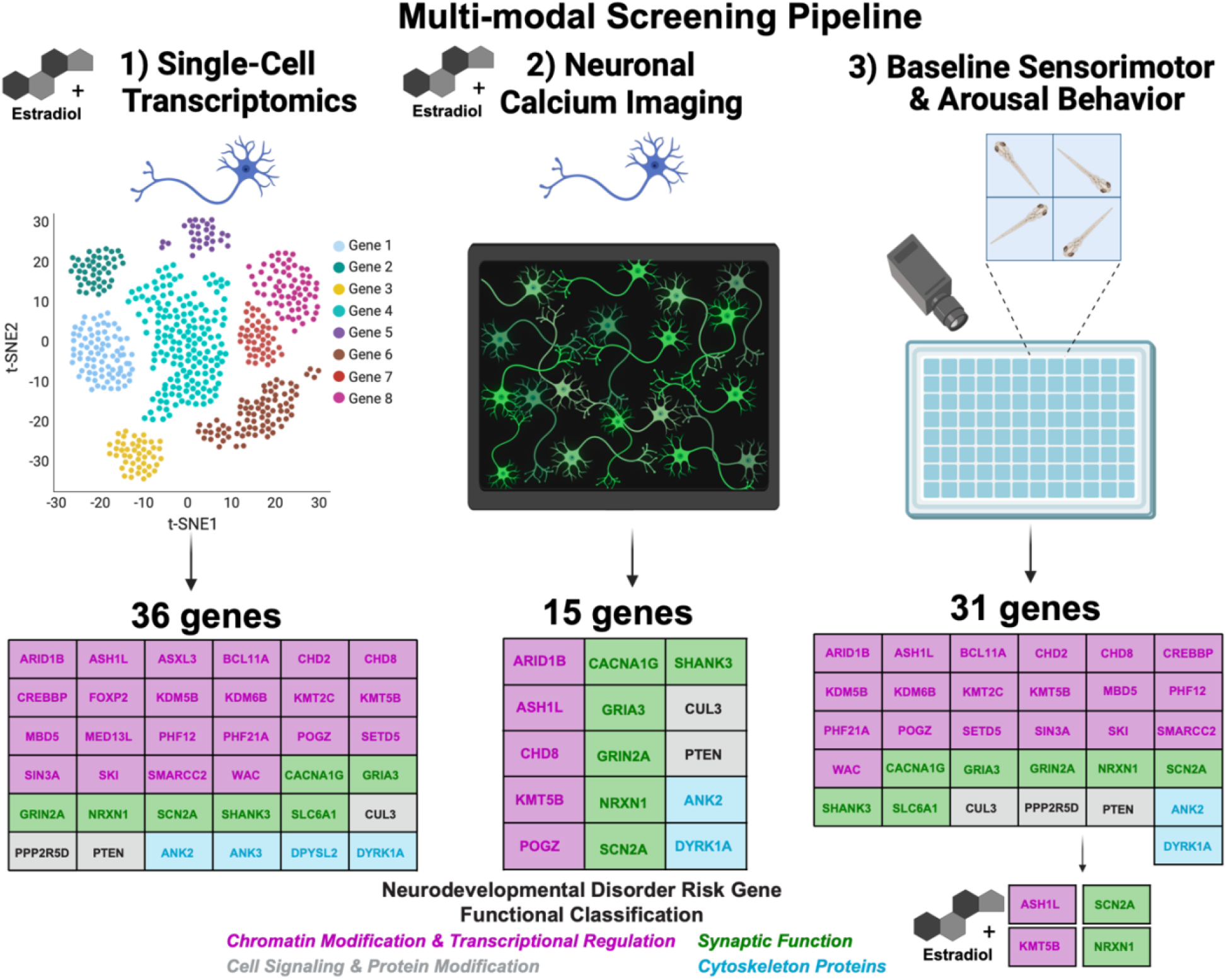
Experimental overview of dual-system, large-scale, multi-modal screening pipeline assessing the impact of 17β-estradiol-x-neurodevelopmental disorder risk genes. The effect of estradiol treatment and loss-of-function mutations in neurodevelopmental disorder (NDD) risk genes is measured across three modalities: (1) single-cell RNAseq in hiPSC-derived *NGN2*-induced glutamatergic neurons (iGLUTs), (2) neuronal calcium imaging in iGLUTs, and (3) sensorimotor and arousal behavior in larval zebrafish. Modalities include overlapping but independent gene sets. ASD/NDD target genes encompass myriad functions, including gene expression regulation (pink), neuronal function and communication (green), cellular signaling and protein modification (gray), and cytoskeleton (blue).

Using pooled and arrayed CRISPR-based techniques, we investigate context-specific and gene knockout-dependent effects of estradiol across cell types and developmental stages, probing *in vitro* gene expression (**Fig. 1-1**: 36 genes) and neuronal activity in human induced glutamatergic neurons (iGLUTs) (**Fig. 1-2**: 15 genes), and *in vivo* sensorimotor and arousal behavior in larval zebrafish (**Fig. 1-3**: 31 genes at baseline, four genes treated with estradiol). In one top gene, *scn1lab*, we assess estradiol effects on neurodevelopmental phenotypes using *in vivo* bulk RNAseq, brain activity mapping, and seizure assays. While ASD/NDD gene knockouts displayed a range of convergent and divergent rescue effects following estradiol exposure, *ASH1L* and *SCN2A* were the top ASD/NDD gene knockouts most amenable to estradiol rescue across modalities. To our knowledge, this study is the first to conduct a systematic, large-scale gene-x-estradiol screen with multiple functional modalities across conserved species and using multiple human donors. Altogether, we systematically assess the interaction between estradiol and mutations in diverse ASD/NDD genes to understand mechanisms and contributions of sex hormones to NDDs.

**Figure 2:**
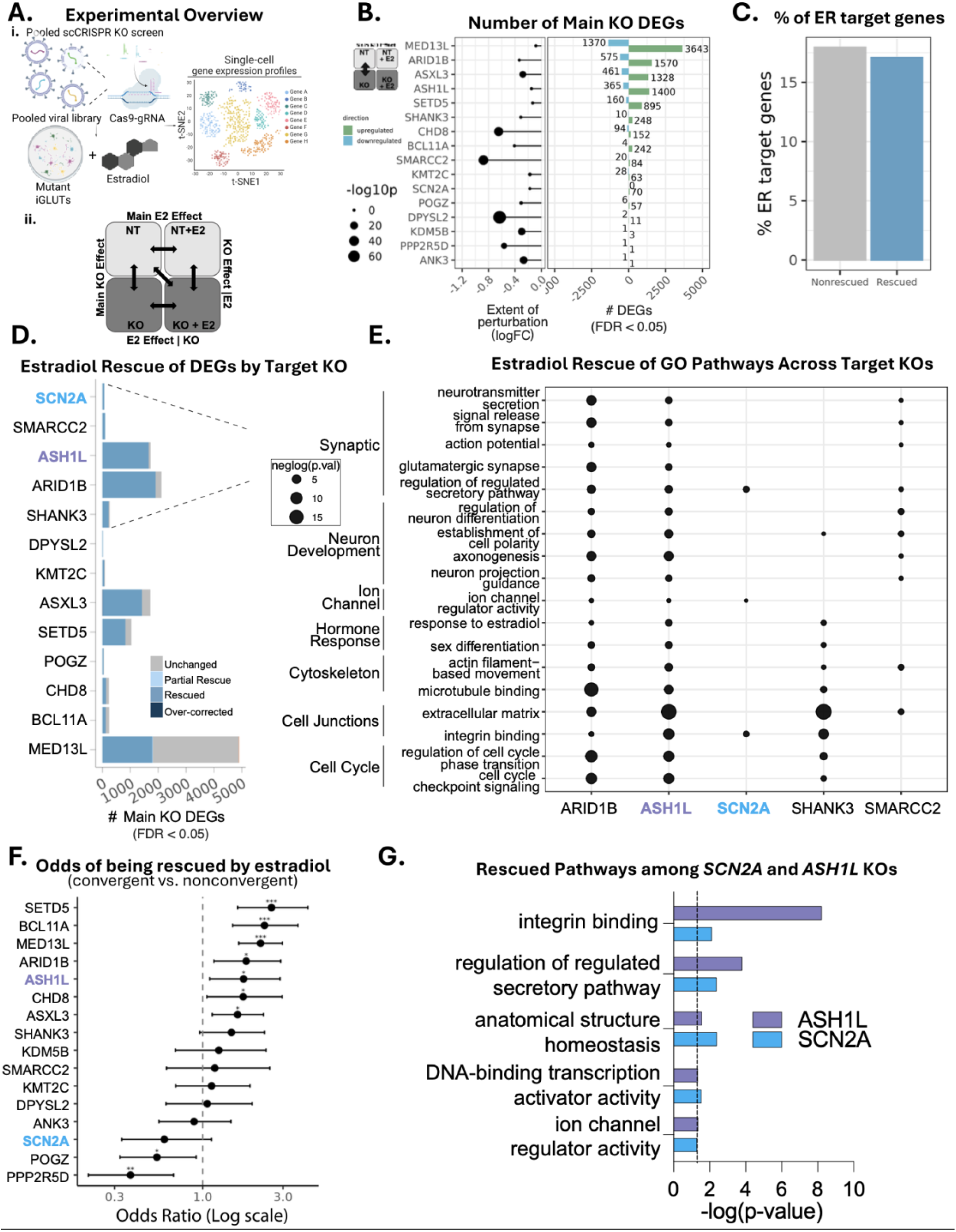
Estradiol rescues dysregulated DEGs and synaptic pathways in iGLUTs. **A.** Schematic of experimental design and different effect comparisons. Thirty-six NDD risk genes are targeted using pooled single-cell CRISPR-knockout screening in hiPSC- derived NGN2-glutamatergic neurons (iGLUTs) with and without 24hr exposure to 100nM estradiol. Each gene was targeted with 3-4 gRNAs **(i).** Various effect comparisons were performed among the treated and untreated groups **(ii).** Main KO effect measures KO+DMSO vs Non-targeting control (NT)+DMSO. Main E2 effect measures NT+E2 vs NT+DMSO. KO effect within E2 measures KO+E2 vs NT+E2. E2 effect within KO measures KO+E2 vs KO+DMSO. E2 and KO joint effect measures KO+E2 vs NT+DMSO. **B.** Extent of perturbation per target NDD gene (left, dot plot). Circle size indicates negative log10(p-value). Stem indicates the log fold change of the knockout +DMSO vs non- targeting control + DMSO. KOs were called based on enrichment of gRNA for target NDD genes and downregulation of target KO gene expression. Number of differentially expressed genes in the main KO effect are shown in the bar plot (FDR <0.05). **C.** Percentage of ER target genes within estradiol rescued genes (blue) or nonrescued genes (gray). **D.** Effect of estradiol rescue on DEGs across NDD risk genes in iGLUTs. Impact of estradiol on number of main KO DEGs is classified into categories of *unchanged, partial rescue, rescue,* and *over-corrected.* For detailed description of degree of rescue thresholds, see Methods and SI Fig 16. Target genes are ranked from highest to lowest percentage of DEGs that are classified as rescued (FDR <0.05). **E.** Gene Ontology (GO) biological pathways rescued by estradiol across the top 5 ranked target KOs. Dot size indicates the –log10(p-value) of the gene set enrichment significance of E2 effect within KO. **F.** Odds of convergent genes being rescued by E2 compared to nonconvergent genes, tested across NDD KO. Fisher’s exact test, ***FDR<0.001, **FDR<0.01, *FDR<0.05. **G.** Select GO terms indicating estradiol rescue among *ASH1L* and *SCN2A* KO in iGLUTs.

**Figure 3:**
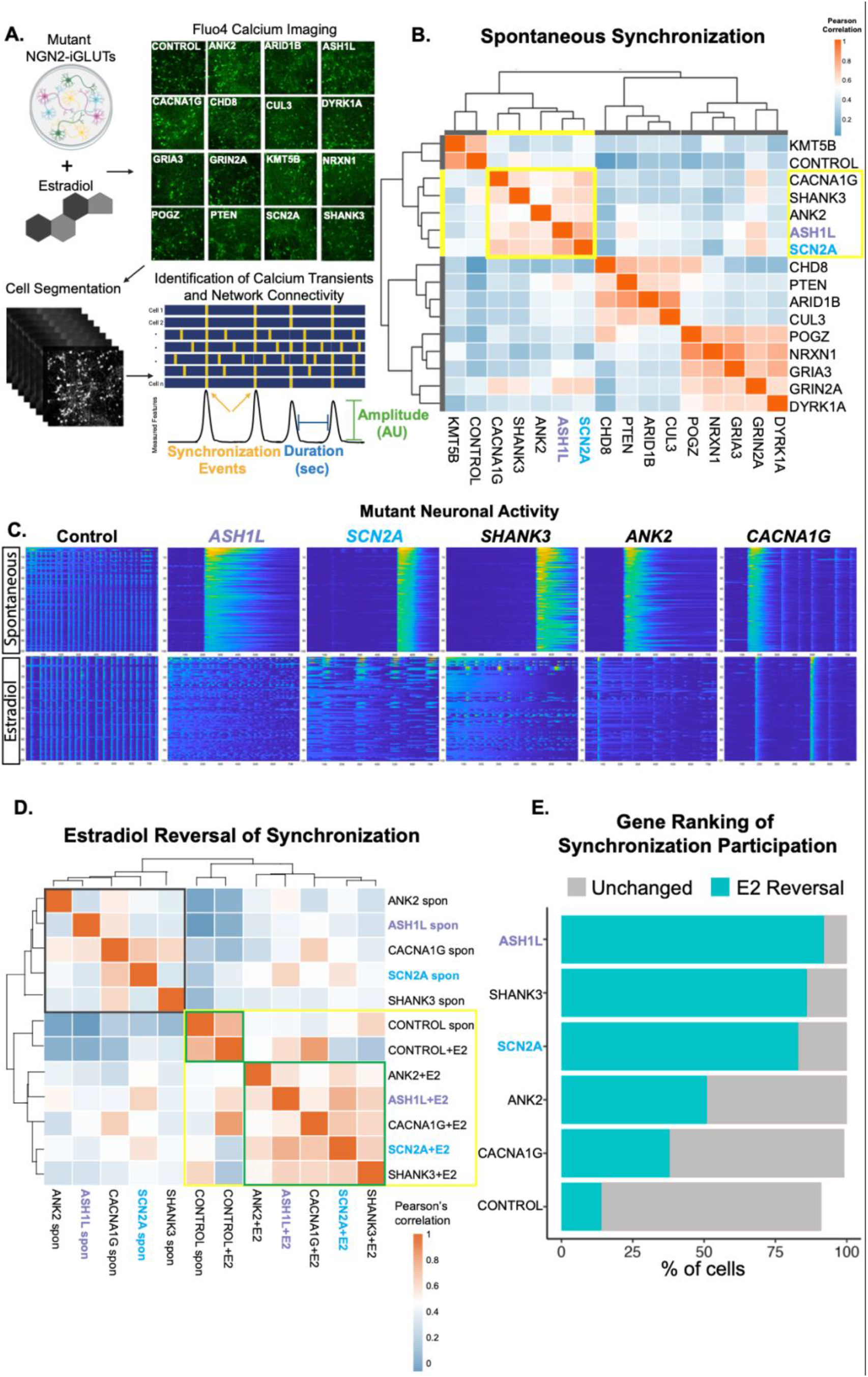
Estradiol rescues hypersynchronous firing pattern across five NDD mutants. **A.** Experimental workflow to assess neuronal activity. Neuronal activity after acute immediate administration of 100nM estradiol is measured in 15 NDD risk gene knockouts in iGLUTs using Fluo4 calcium imaging with confocal microscopy. Individual cells are identified using customized pixel-wised cell segmentation analysis. Duration and amplitude of calcium transients as well as network synchronization events are measured. **B.** Correlation analysis across spontaneous network synchronization identifies 4 sub- clusters of NDD risk genes with related phenotypes. Cluster 1: *KMT5B* and *non-targeting control (NT).* Cluster 2: *CACNA1G, SHANK3, ANK2, ASH1L, SCN2A.* Cluster 3: *CHD8, PTEN, ARID1B, CUL3.* Cluster 4: *POGZ, NRXN1, GRIA3, GRIN2A, DYRK1A*. Yellow box indicates cluster 2 with a distinct hypersynchronous firing phenotype. **C.** Raster plots of spontaneous baseline neuronal activity (top) and estradiol treated (bottom) in non-targeting control, *ASH1L, SCN2A, SHANK3, ANK2, and CACNA1G.* Each row of the raster plot is a cell, and each column is a frame. Cells were imaged for 1 minute (750 frames). **D.** Correlation analysis across spontaneous and estradiol-treated network synchronization. Two distinct clusters are formed. Cluster 1: Spontaneous *ANK2*, *ASH1L, CACNA1G, SCN2A, and SHANK3.* Cluster 2: Spontaneous non-targeting control, Estradiol-treated Non-targeting control, Estradiol-treated *ANK2, ASH1L, CACNA1G, SCN2A, SHANK3.* **E.** Ranking of risk gene knockouts by percent of active neurons that contribute to synchronization pattern that are reversed by estradiol treatment.

## RESULTS

### Estradiol rescues transcriptional effects of ASD/NDD risk genes

Towards resolving shared and distinct transcriptomic effects of estradiol across ASD/NDD risk genes, we prioritized 36 ASD/NDD genes with robust LOF effects (**Fig. 1-1, SI Table 1**), focused on 22 gene expression regulators (e.g., epigenetic modifiers, chromatin regulators and transcription factors: *ARID1B, ASH1L, ASXL3, BCL11A, CHD2, CHD8, CREBBP, FOXP2, KDM5B, KDM6B, KMT2C, KMT5B (SUV420H1), MBD5, MED13L, PHF12, PHF21A, POGZ, SETD5, SIN3A, SKI, SMARCC2, WAC*), but also including 7 neuronal communication genes (e.g., synaptic and ion channel genes: *CACNA1G, GRIA3, GRIN2A, NRXN1, SCN2A, SHANK3, SLC6A1)*, 3 cell signaling genes (*CUL3, PPP2R5D, PTEN)*, and 4 cytoskeleton genes (*ANK2, ANK3, DPYSL2, DYRK1A)*. Gene selection was not based on sex-differential penetrance; many have similar penetrance in males and females^42^ (**SI Table 1**). The lentiviral CRISPR-KO library was generated from pre-validated gRNAs (three to four gRNAs per gene; **SI Table 2, SI Table 3**), 29 of which were included and validated in our previous study^43^.

We applied a pooled CRISPR-knockout (KO) approach (**Fig. 2Ai)** in hiPSC-derived NPCs^44^ (derived from two controls: one male and one female) (**SI** Fig. 1A) and functionally mature iGLUTs^45^ (derived from one control: male)^46^ (**Fig. 2, SI Fig. 1A**) treated with estradiol (E2, 100nM, 24 hours, receptor expression **SI Fig. 2**, dose response assays **SI Fig 3A**) or vehicle control (DMSO). Direct detection of sgRNAs was coupled with single-cell RNA sequencing (ECCITE-seq^47^) to compare effects across cell types and E2 contexts; gRNAs were validated (**SI Fig. 4**). Altogether, we resolved 39,278 cells across 16 ASD/NDD KOs (and scramble controls) after filtering and quality control: 7,067 vehicle NPC, 5,553 E2 NPC, 14,471 vehicle iGLUT, and 12,187 E2 iGLUT (**SI Fig. 1B,C, SI Table 4**). We combined target-gene depletion filtering with weighted-nearest network (WNN) analysis to identify cells with successful perturbations, assigning clusters to specific knockouts based on enrichment of gRNA and gene expression profiles. This strategy yielded a median of 234 (iGLUT) and 198 (NPC) cells per gRNA and 39 (iGLUT) and 28 (NPC) gRNAs identified in >75 cells. On average, CRISPR-KO led to 22% (iGLUT) and 26% (NPC) target gene depletion, which, owing to the heterogeneity of nonsense mediated decay within cell populations^48^, likely represents an underestimate of true target gene perturbation effects^49^.

Transcriptomic signatures were compared to non-targeting scramble control (NT) clusters across multiple axes to resolve knockout effects, estradiol effects, and knockout-x-estradiol interact effects (**Fig. 2Aii).** At baseline, each ASD/NDD resulted in shared and distinct differentially expressed genes (DEGs, FDR <0.05), not determined by the extent of perturbation of each target gene (**Fig. 2B, SI Fig. 1D**). For the ten ASD/NDD genes successfully targeted by two or more gRNAs, we observed robust correlations of DEGs (Pearson correlation 0.66-0.87) (**SI Fig 4B**).

For all significant differences between KO and control (p<0.05), we define *exacerbated* as a perturbation in the same direction that is more significant, *unchanged* as no change in the direction or significance of effect, *partial rescue* as a weaker perturbation that is less significant in the original direction, *rescued* as the loss of any significant difference between mutant and control, and *over-corrected* as a difference in the opposite direction of effect of the original phenotype (see **Methods and SI Fig 16** for more details). The impact of estradiol on DEGs (FDR<0.05) and biological pathways (FDR <0.05) was assessed across target KOs. Estradiol rescued DEGs across 13 resolved KOs, albeit with heterogeneity in degree of rescue (iGLUT FDR <0.05, ranked by percentage of DEGs rescued, **Fig. 2D**, NPC nominal p-value <0.05, **SI Fig. 1E**). Across this and all subsequent analyses, we highlight two genes with robust multimodal estradiol rescue effects: *ASH1L* and *SCN2A*. As representative examples of highly rescued genes, for each degree of rescue we compared the effect sizes of the main KO effect to the estradiol treated KO effect (**SI Fig. 5, top panel**) and the main KO effect to the estradiol effect in KO (**SI Fig. 5, bottom panel**). The effect sizes of the main KO genes were ameliorated when treated with estradiol; KO effects and estradiol effects were negatively correlated.

We previously examined cell-type-specific downstream impacts between NDD/ASD risk genes at baseline, reporting that points of convergence were greatest in mature iGLUTs, where they broadly targeted synaptic and epigenetic pathways and networks^59^. Unexpectedly, rescued DEGs in the current dataset were neither enriched for nor more likely to be ER target genes compared to nonrescued DEGs, suggesting that they are mediated, at least in part, by non-genomic signaling pathways (**Fig. 2C, SI Fig. 6A, 6C-G**). Convergent DEGs, defined as those with significant meta-analysis FDR (FDR < 0.05), non-significant Cochran’s heterogeneity Q test p (p > 0.05), and shared direction across all 16 resolved KOs, were more likely to be ameliorated by estradiol vs nonconvergent DEGs, defined as those that did not meet the convergence threshold, in 7 gene targets, but not more likely to be an ER target, further suggesting non-genomic mechanisms (Fisher’s exact test, ***FDR<0.001, **FDR<0.01, *FDR<0.05) **(Fig. 2F, SI Fig. 6B)**.

All Gene Ontology (GO) pathways^50,51^ that were significant in the KO main effect (nominal p-value <0.05) were compared to the estradiol effect within KO. Indeed, in iGLUTs we resolved shared dysregulated pathways that estradiol rescued, notably related to synaptic transmission, neuronal development, cytoskeleton organization, cellular junctions, cell cycle, hormone response, and ion channel activity across the top 5 KOs. (**Fig. 2E, SI Table 5)**. Here we note that for *ASH1L* and *SCN2A*, estradiol rescued pathways were associated with integrins, structural homeostasis, and transcriptional activity in iGLUTs (**Fig. 2G**). For 8 KOs in NPCs, estradiol rescued pathways related to transcription, translation, membrane transport activity, mitochondrial energy production, and cell junctions (**SI Fig. 1F-G, 1G, SI Table 6**).

### Rapid reversal of network hypersynchronicity in ASD/NDD neurons by estradiol

To more specifically query non-genomic effects of estradiol in iGLUTs, we combined an arrayed CRISPR-KO approach with calcium imaging (Fluo4) to assess the immediate impact of estradiol exposure (100nM, immediate) on neuronal network activity across 15 genes, balanced to now focus equally on gene expression regulators *(ARID1B, ASH1L, CHD8, KMT5B, POGZ)* and neuronal communication *(CACNA1G, GRIA3, GRIN2A, NRXN1, SCN2A, SHANK3)* genes, and retaining a smaller number of cytoskeletal (*ANK2, DYRK1A)* and cell signaling-protein modification (*CUL3, PTEN*) genes (**Fig. 1-2)**. Because neuronal activity dynamics of iGLUTs change over development, we assayed cell-autonomous spiking of individual neurons (i.e., calcium transient frequency, duration, and amplitude) (30DIV; immature networks) and non-cell-autonomous network activity (i.e., synchronized oscillations and network bursts) (60DIV; mature networks) at distinct timepoints (**Fig. 3A; SI Fig. 7, SI Fig. 8A**).

The 15 ASD/NDD genes showed heterogeneous neuronal effects across genes and neurodevelopmental timepoints, with a subset showing robust estradiol reversal. In immature networks (30DIV), estradiol treatment ameliorated spiking frequency in five KOs (*SHANK3, DYRK1A, POGZ, GRIA3,* and *GRIN2A*) and amplitude in six KOs (*DYRK1A, POGZ, GRIA3, GRIN2A, CACNA1G, SHANK3)* (**SI Fig. 7)**. In mature networks (60DIV), four distinct network synchronization clusters were resolved at baseline (**Fig. 3B**; **SI Fig. 8B-E**): cluster 1 (control) displayed typical network synchronization; cluster 2 (long-lasting synchronization events) revealed a sustained burst (*ASH1L, SCN2A, SHANK3, ANK2, CACNA1G*); cluster 3 (network hypoactivity) lacked network oscillation: *ARID1B, CUL3, CHD8,* and *PTEN);* and cluster 4 (intermediate) exhibited sparse and smaller synchronization events (*NRXN1, GRIA3, POGZ, GRIN2A, DYRK1A)* (**Fig. 3B, 3C, SI Fig. 8B-E**). KO neurons treated with estradiol clustered more closely with controls (**Fig. 3D**), but cluster 2 revealed the most robust estradiol reversal, with rapid amelioration of the hypersynchronicity and restored oscillatory-like activity patterns immediately after estradiol treatment (**Fig. 3C**), with the extent of rescue ranked based on the percentage of synchronized cells (**Fig. 3E**). Overall, we identified a subset of ASD/NDD risk genes that exhibited hypersynchronous firing activity that were immediately reversed by estradiol treatment (*ASH1L, SCN2A, SHANK3, ANK2,* and *CACNA1G),* consistent with non-genomic mechanisms.

### *In vivo* rescue of neurodevelopmental and sleep-related dysregulation by estradiol

We evaluated baseline sensorimotor and arousal behavior across 31 zebrafish ortholog(s) of ASD/NDD genes (CRISPant F0 mosaic knockouts^52^ or stable mutant lines^38,43,53–55^), including 19 gene expression regulators (*ARID1B, ASH1L, BCL11A, CHD2, CHD8, CREBBP, KDM5B, KDM6B, KMT2C, KMT5B, MDB5, PHF12, PHF21A, POGZ, SETD5, SIN3A, SKI, SMARCC2, WAC*), 7 neuronal communication genes (*CACNA1G, GRIA3, GRIN2A, NRXN1, SCN2A, SHANK3, SLC6A1)*, 3 cell signaling-protein modification genes (*CUL3, PPP2R5D, PTEN)*, and 2 cytoskeletal genes (*ANK2, DYRK1A)* (**Fig. 1-3; SI Tables 8-11)**. In zebrafish, *scn1lab* is an ortholog of human *SCN1A* and *SCN2A*.

Hierarchical clustering of quantitative behavioral profiling (via high-throughput automated video tracking) across 24 visual-startle (VSR)^38,56^ and sleep-wake parameters^57,58^ (**Fig. 4A, SI Table 7**) revealed seven behavioral clusters at baseline: group 1 (*wacab, crebbpab, arid1b, phf21aab*), group 2 (*cacna1g, ptenab, nrxn1ab, scn1lab, ash1l, chd8*), group 3 (*grin2aab, chd2, skiab, phf12ab*), group 4 (*kdm5ab, ppp2r5d, bcl11ab, mbd5*), group 5 (*ank2ab, kmt2cab, kmt5b*), group 6 (*slc6a1ab, kdm6bab, dyrk1aab, sin3ab*), group 7 (*smarcc2, setd5, gria3ab, pogzab, shank3ab, cul3ab*) (**Fig. 4C; SI Fig. 9A)**. We leveraged our pharmaco-behavioral database^59^ to identify drugs predicted to (i) mimic or positively correlate with the mutant phenotype or (ii) oppose or anti-correlate with the mutant phenotype (**Fig. 4B**), asking if drugs targeting genomic (ERα and ERβ) or nongenomic (GPER1) mechanisms were enriched at the gene (**Fig. 4D, SI Table 13**) or cluster (**SI Fig. 9B**) level. Target genes for which there was no estradiol enrichment (*cacna1g, pten, nrxn1)* are not shown in the enrichment plot. Behavioral phenotypes of ASD/NDD gene mutants in cluster 2, particularly *ash1l* and *scn1lab*, showed the strongest anti-correlation with estrogenic compounds (anti-correlation <-0.5, p<0.05, t-statistic) (**SI Table 14**); in contrast, ASD/NDD gene mutants in other clusters (*smarcc2, setd5*, *gria3ab, shank3ab,* and *kmt5b*) displayed phenotypes that highly correlated with the effects of estradiol treatment in wild-type fish (correlation >0.5, p<0.05, t-statistic). Consistent with a plausible role for genomic signaling mechanisms, protein-protein interaction (PPI) networks (STRING^60^) for the top predicted estradiol responder genes, *ash1l* (**Fig 4Ei, SI Fig. 9C**) and *scn1lab*^59^, included ERα, ERβ, and GPER as anti-correlating drug targets and molecular pathways (GO^50,51^) enriched for nuclear transcription, hormone-mediated signaling, and estrogen signaling pathways (**Fig. 4Eii**).

**Figure 4:**
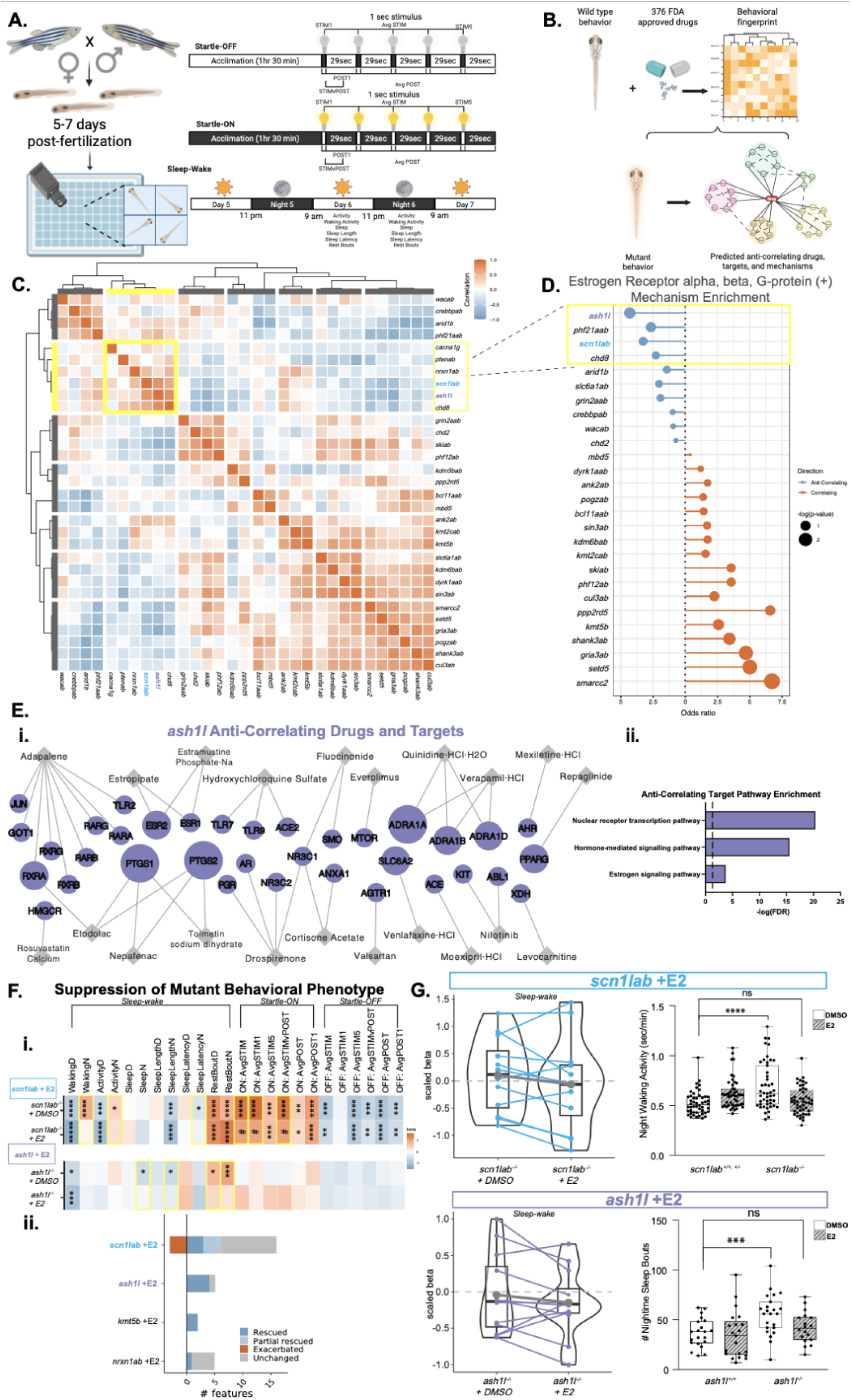
Estradiol receptor mechanisms predict estradiol rescue of arousal and sensorimotor phenotypes in ash1l and scn1lab mutants. **A.** Experimental setup of larval zebrafish behavior assays. Larval behavior is measured through automated video- tracking. *Visual-startle (VSR)*: Larvae acclimate and are exposed to five 1-second flashes of lights-off (Startle-OFF) or -on (Startle-ON) stimuli at 29-second intervals at 5 days post- fertilization (dpf). *Sleep-wake*: Larvae are exposed to a 14h:10h light:dark cycle from 5- 7dpf. Visual-startle and sleep-wake parameters are depicted. **B.** Schematic of pharmaco- behavioral profiling of 376 FDA-approved compounds in wild-type zebrafish to predict drugs and mechanisms that anti-correlate (oppose) or correlate (mimic) with mutant behavioral phenotypes. **C.** Correlation analysis across 24 visual-startle and sleep-wake behavioral parameters identifies 7 sub-clusters of NDD risk gene mutants with related phenotypes. Cluster 1: *wacab, crebbpab, arid1b, phf21aab.* Cluster 2: *cacna1g, ptenab, nrxn1ab, scn1lab, ash1l, chd8.* Cluster 3: *grin2aab, chd2, skiab, phf12ab.* Cluster 4: *kdm5bab, ppp2rd5, bcl11aab, mbd5.* Cluster 5: *ank2ab, kmt2cab, kmt5b.* Cluster 6: *slc6a1ab, kdm6bab, dyrk1aab, sin3ab.* Cluster 7: *smarcc2, setd5, gria3ab, pogzab, shank3ab, cul3ab.* F0 and stable line mutation sequences are found in **SI Tables 8-9.** Yellow box highlights cluster 2, which includes genes from distinct functional classifications. **D.** Molecular targets *estrogen receptor (ER) alpha, beta, and G-protein* are significantly enriched among anti-correlating (blue, correlation <-0.5, p<0.05, t- statistic) and correlating drugs (orange, correlation >0.5, p<0.05, t-statistic) drugs across NDD gene mutants. Mutants for which there is no ER enrichment are not shown. Circle size indicates the −log10(p-value), calculated using Fisher’s exact test. Stem length indicates the odds ratio for each mutant. NDD genes are ranked by circle size and stem length. Enrichment of estrogen receptor alpha, beta, and G-protein and p-values for each gene are shown in **SI Table 13. E.** Protein-Protein Interaction (PPI) network among the targets (purple circles) of anti-correlating drugs (gray diamonds) (correlation <-0.5, p<0.05, t-statistic) for *ash1l* mutant based on the STRING database **(i).** Node size of proteins represents the number of drugs targeting each protein. GO pathways enriched among the drug targets are indicated **(ii). F.** Heatmap of empirical beta values for each behavioral parameter for *scn1lab* and *ash1l* mutants between the effect of the mutant alone (top row, HOM DMSO vs WT DMSO) and for the effect of the interaction of the mutant estradiol treatment (24 h, 20 µM) compared to WT vehicle (HOM E2 vs WT DMSO), showing relative suppression of behavioral features. Asterisks (*) indicate behavioral features that are significantly different in indicated comparison group (p * <0.05, ** <0.01, ***<0.001, ****<0.0001, LMM). Yellow boxes indicate parameters that were rescued (significantly different in HOM DMSO vs WT DMSO and no longer significant in HOM E2 vs WT DMSO) or partially rescued (significantly different in HOM DMSO vs WT DMSO and reduced in HOM E2 vs WT DMSO). Pound sign (#) indicates feature is partially rescued **(i).** For behavioral features that are significantly different in the HOM DMSO vs WT DMSO p-value ≤ 0.05, we characterized the effect of the HOM-x-E2 for each feature as *exacerbated, unchanged, partial rescue, rescue,* and *over-corrected* **(ii).** For detailed description of the thresholds for each classification, see Methods and **SI Fig 16**. Mutants are ranked by the number of features that are *rescued* or *partially rescued*. **G.** For *scn1lab* and *ash1l* mutants, violin plots show comparison of the magnitude of effect (distribution of scaled beta scores) on sleep-wake behavioral parameters between the HOM DMSO vs WT DMSO (left) and the HOM E2 vs WT DMSO (right). Box and whisker plots show representative parameters that were rescued for each mutant. Increased nighttime waking activity and increased number of sleep bouts were rescued by estradiol in *scn1lab* and *ash1l* mutants, respectively (Unpaired t-test with Welch’s correction, **** p < 0.0001, *** p = 0.0010. *scn1lab* n = 48/group; *ash1l* WT DMSO n=21, WT E2 n=21, HOM DMSO n=23, HOM E2 n=18).

The rescue effect of estradiol was experimentally evaluated across *ash1l* and *scn1lab* mutants (predicted reversal), *kmt5b* mutants (predicted mimic), and *nrxn1ab* mutants (no predicted effect) (**Fig. 4F**). The visual-startle stimuli and sleep-wake activity of 5-7 dpf homozygous mutants or control larvae was measured following treatment with estradiol (20 µM, 24 hours, dose response assay **SI Fig. 3B-3C**) or vehicle control (DMSO). The phenotypic impact of mutant-x-estradiol combinations was ranked based upon the total number of features partially rescued and rescued; consistent with predictions, *scn1lab* and *ash1l* mutants showed greatest estradiol rescue, followed by *kmt5b* mutants, and finally *nrxn1ab* mutants (**Fig. 4Fii**). Of note, immediate estradiol treatment (presumably via non-genomic mechanisms) resulted in similar gene-specific rescues (**SI Fig. 10, SI Table 12**), although the 24-hour pre-treatment condition showed a stronger magnitude of rescue. Estradiol rescued specific sleep-wake parameters in *scn1lab* and *ash1l* mutants, ameliorating increased nighttime activity and sleep bouts, respectively (**Fig. 4G**).

Altogether, we describe congruence of gene-specific baseline behavioral phenotypes, predicted responsivity to estrogenic compounds, and experimental rescue, particularly related to hyperexcitability behavioral phenotypes associated with select NDD genes.

### Estradiol rescues transcriptional and network activity dysregulation in *SCN2A* **neurons and zebrafish *scn1lab* mutants**

For *SCN2A,* a top estradiol responsive ASD/NDD gene, with a particularly pronounced cross-species hyperexcitability phenotype at baseline61-63, we sought to uncover *in vivo* transcriptomic and cellular mechanisms linked to estradiol rescue.

Background-matched *scn1lab* homozygous mutant and wild-type zebrafish larvae were treated with estradiol (20 µM dissolved in DMSO, 24 hours from 5-6 dpf) or vehicle (DMSO) followed by whole-brain bulk RNAseq (**Fig. 5A**). Consistent with our scRNAseq data (**Fig. 2D-2E**), the significant DEGs (nominal p <0.1) and biological pathways (GO^50,51^, FDR <0.05) dysregulated in *scn1lab* homozygous mutant brains but rescued with estradiol treatment include enrichments for synaptic, neuron development, and the cell cycle, with additional enrichment for mitochondrial-mediated energy-related pathways and ion channel activity. (**Fig. 5B-C, SI Fig. 11, SI Tables 15-16)**. Yet, enrichment for estrogenic pathways suggests that transcriptomic differences might be mediated by estrogen-dependent genomic signaling.

**Figure 5:**
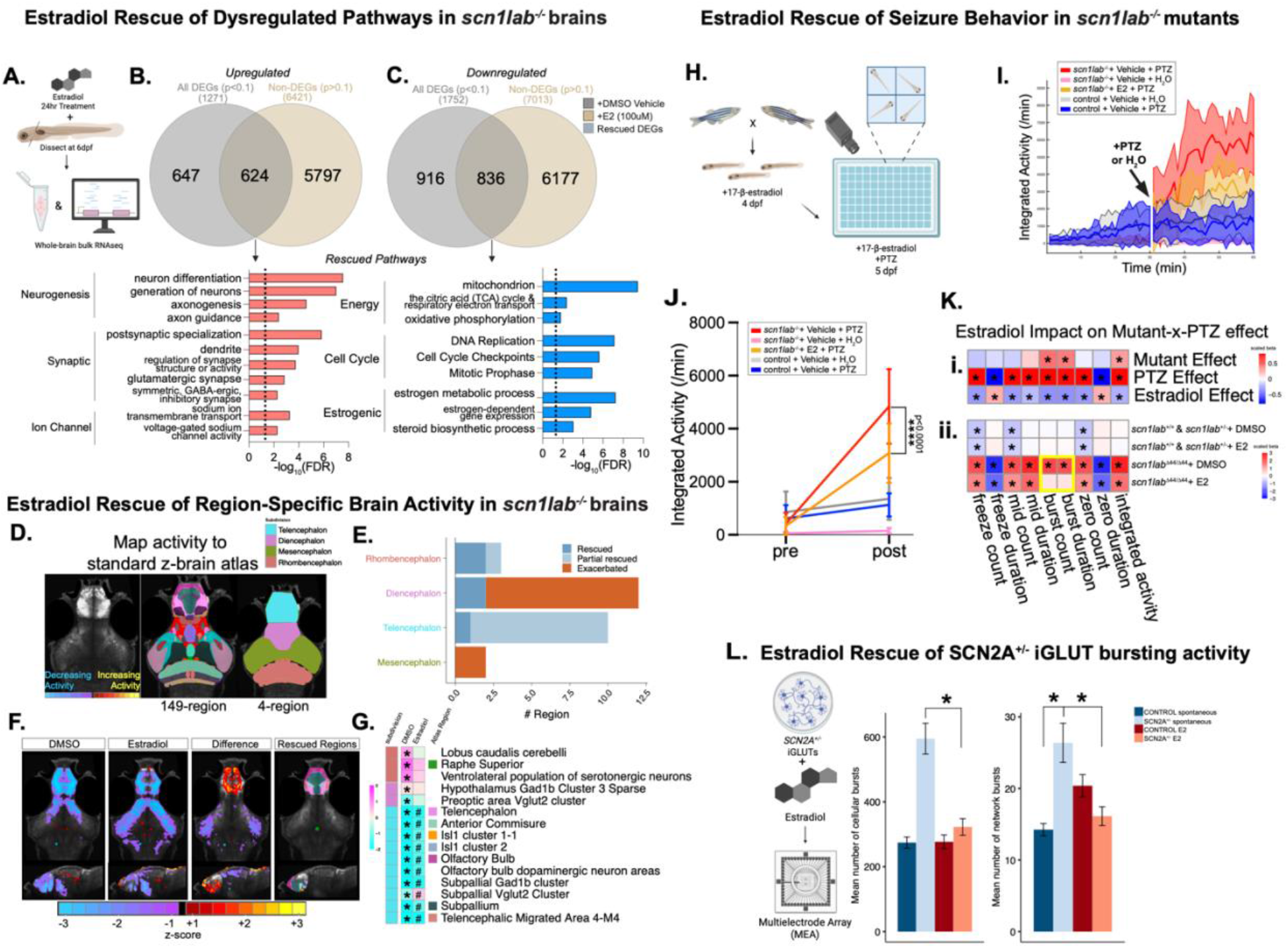
Estradiol rescues transcriptional and neuronal network activity dysregulation in *scn1lab*/*SCN2A* mutants. **A.** Schematic of experimental workflow to assess impact of estradiol on baseline gene expression in *scn1lab^-/-^* larvae. **B-C**. Venn diagrams showing the overlap of upregulated (**B**) or downregulated (**C**) DEGs (p <0.1) in *scn1lab^-/-^* + DMSO and non-DEGs (p >0.1) in *scn1lab^-/-^* + estradiol (E2), representing rescued DEGs. Venn diagrams made in InteractiVenn^155^. Gene Ontology (GO)^50^ pathways enriched among upregulated (red) and downregulated (blue) rescued DEGs are shown below the Venn diagrams. Dashed line represents FDR <0.05. For the full list of rescued pathways, see SI Table 16. **D.** Schematic of method to map pERK/tERK activity to a 149 and 4 region z-brain atlas. **(E)** Estradiol rescued regions by 4-region atlas. **(F)** Voxel-wise Z score normalized pERK/tERK values representing brain activity differences in *scn1lab^Δ44/Δ44^* + DMSO or E2 compared to DMSO-treated *scn1lab^+/+^* larvae, the difference between the two conditions from a representative experiment, and rescued regions from the 149-region atlas in the three experiment meta-analysis. For visualizations of all three replicate experiments, see SI Fig. 13. **(G)** Heat map showing regions from the 149-region atlas with significant differences in activity (pERK/tERK) in *scn1lab^Δ44/Δ44^* + DMSO vs *scn1lab^+/+^* + DMSO brains (p <0.05, LMM meta-analysis) that are rescued in *scn1lab^Δ44/Δ44^* + E2 (p >0.05, LMM meta-analysis) or partially rescued (for degree of rescue threshold criteria, see Methods and **SI Fig. 16**). Each rectangle represents the scaled beta from LMM comparing brain activity in *scn1lab^Δ44/Δ44^* + E2 or DMSO to *scn1lab^+/+^* + DMSO (magenta, higher than *scn1lab^+/+^* + DMSO; cyan, lower than *scn1lab^+/+^* + DMSO). Colors shown next to region names refer to the “Rescued Regions” image in panel F. Asterisks (*) indicate regions that are significant in the DMSO condition. Pound sign (#) indicates regions that are partially rescued in E2 condition. Empty box in the E2 condition indicates full rescue. **H.** Left: Schematic of experimental workflow to assess impact of estradiol on induced seizure activity in *scn1lab^-/-^* larvae. **I.** Pentylenetetrazole (PTZ)-induced seizure activity, with and without 24 h exposure to 20 µM estradiol, is measured in *scn1lab^-/-^* larval zebrafish mutants using video-tracking. Representative activity tracings of control and *scn1lab*^-/-^ larvae at 5dpf before and after exposure to DMSO vehicle or 20 µM estradiol and 10 mM PTZ or water. Ribbon shows +/- SEM. **J.** Average activity 30 min before (pre) and after (post) the addition of 10mM PTZ or water to control and *scn1lab*^-/-^ larvae before and after exposure to DMSO vehicle or 20 µM estradiol. *scn1lab*^-/-^ + E2 + PTZ larvae show significantly decreased activity compared to *scn1lab*^-/-^ + vehicle + PTZ larvae (Unpaired t-test with Welch’s correction on post values, ****p<0.0001; n=24/group). **K.** Interaction models assessing genotype-x- estradiol-x-PTZ effect, comparing post- vs pre-PTZ measurements. Dissection of the mutant effect (HOM vs control), estradiol effect (Estradiol vs DMSO), and PTZ effect (PTZ vs water) (i). Three-way interaction model to estimate the PTZ effect among genotype-x- estradiol groups of larvae (ii). **L.** Left: Schematic of experimental workflow to assess impact of estradiol on neuronal busting activity in CRISPR-engineered *SCN2A^+/-^ iGLUTs*. Right: Cellular and network bursting activity, with and without exposure to acute 100nM estradiol, is measured in *SCN2A^+/-^* NGN2-iGLUTs using multielectrode array. Bar graphs show mean number of cellular bursts and network bursts in controls and *SCN2A^+/-^* iGLUTs. Unpaired t-test, cellular bursts *p=0.0415, network bursts *SCN2A^+/-^* spontaneous vs *SCN2A^+/-^* E2 *p=0.0127, *SCN2A^+/-^* spontaneous vs WT spontaneous *p=0.0450.

To determine the effect of estradiol on circuit-level dysregulation *in vivo*, we assessed *scn1lab* mutant zebrafish baseline brain activity and induced seizures following estradiol exposure.

First, whole-brain activity mapping was performed in background-matched wild-type and homozygous *scn1lab* mutants (6 dpf) by immunostaining for phosphorylated ERK (pERK), which responds quickly to influx of calcium after neuronal activity, relative to total ERK (tERK)^64^ (**Fig. 5D**). The pERK/tERK ratio is a well-established, widely used proxy for neuronal activity that has been robustly validated by our group and others for baseline and stimulus-driven activity^38,55,59,65^. Although estradiol has been shown to induce rapid activation of MAPK signaling^66^, there was no overall difference in tERK intensity alone in wild-type fish post-estradiol treatment across three independent experiments (**SI Fig. 12, SI Table 17**). Estradiol treatment (20 µM, 24 hours) showed partial or full rescue of several regions in the telencephalon, rhombencephalon and diencephalon (**Fig. 5E-F, SI Fig. 13**). Mapping brain activity to a more detailed 149-region neuroanatomical atlas highlighted bidirectional regional rescue activity of estradiol, including hypoactivity in telencephalic regions (e.g., subpallial Vglut2 cluster) and the preoptic area Vglut cluster, and hyperactivity in rhombencephalic regions (e.g., raphe superior) and hypothalamus Gad1b cluster 3 (**Fig. 5G, SI Table 18)**. Reported expression of estrogen receptors (*esr2b* and *gper1*) partially mirrors these results. Whereas *esr1* is not expressed in the brain at larval stages and *esr2a* is not spatially restricted, *esr2b* is specifically expressed in ventral telencephalon, anterior diencephalon/preoptic area, and hypothalamus during larval stages^67,68^, as is *gper1* in the olfactory area, preoptic area, ventral hypothalamus, and pituitary at 48-50 hpf^69^. *esr1, esr2a,* and *esr2b* are expressed in the preoptic area and mediobasal hypothalamus in the brains of adult female zebrafish, where their expression patterns partially overlap^70^.

Second, induced seizure activity was compared across wild-type, heterozygous, and homozygous *scn1lab* mutant larvae following exposure to the GABA-A receptor antagonist pentylenetetrazol (PTZ) (10 mM) to induce seizures^71^ (i.e., rapid burst-like increases in activity and circling motion, followed by inactivity) (**Fig. 5H**). Estradiol treatment (20 µM, 24 hours from 5-6 dpf) significantly decreased PTZ-induced seizures in homozygous *scn1lab* mutants (**Fig. 5I-J**). A three-way linear mixed model (LMM) dissected effects of genotype and treatment across behaviors (**Fig. 5Ki**), revealing that estradiol treatment rescues burst count and burst duration of induced seizures in homozygous *scn1lab* mutants (**Fig. 5Kii, SI Table 19**).

Disentangling genomic and non-genomic mechanisms is complex; for example, nongenomic effects impact cellular signaling, protein trafficking and mitochondrial function, but also CREB phosphorylation^72,73^, leading to rapid increases in expression of *Arc* and other genes^74^. We applied selective agonists for ERα (propyl pyrazole triol, PPT, 20 µM), ERβ (diarylpropionitrile, DPN, 20 µM), and GPER1 (G1, 1 µM and 2 µM) to conduct a focused pharmacological manipulation of estrogen receptors in the context of the PTZ seizure assay. When homozygous *scn1lab* mutant zebrafish larvae were exposed to each drug for 24 hours, and then treated with vehicle or PTZ, all three agonists rescued increased burst count and duration (**SI Fig 14**), suggesting that nuclear and membrane estrogen receptors jointly mediate the effect. Estradiol rescue of increased activity was reinforced in CRISPR engineered heterozygous human SCN2A^+/-^ iGLUTs^75^, whereby acute administration of estradiol (100nM, 24 hours) likewise decreased baseline cellular and network bursting activity measured by multielectrode array (MEA) (**Fig. 5L, SI Table 20**).

Altogether, homozygous *scn1lab* mutant zebrafish reveal regional changes in whole-brain activity coupled with increased susceptibility to induced seizure activity, both of which are ameliorated by estradiol treatment. Consistent with our multi-gene transcriptomic, cellular, and behavioral analyses, whole-brain circuit level studies further indicate that estradiol rescues dysregulated network activity in the context of select ASD/NDD genes.

### Enhanced network connectivity and distinct developmental cortical expression patterns of top estradiol-responsive genes

Integration of multi-modal transcriptomic, cellular, and behavioral datasets was performed to identify ASD/NDD genes across modalities that were most receptive to estradiol rescue (**Fig. 6A, SI Table 21**). Mapping Wasserstein barycenters for each modality onto latent space (**Fig. 6B**) revealed three distinct clusters (gene control, estradiol-treated, vehicle-treated) (**Fig. 6C**). Inter-modality distance across treatment conditions revealed iGLUT calcium imaging (neuronal activity) and zebrafish behavior modalities to be more similar within treatment conditions, and single-cell transcriptomics to be most distinct, both within and across conditions (**Fig. 6D**).

**Figure 6:**
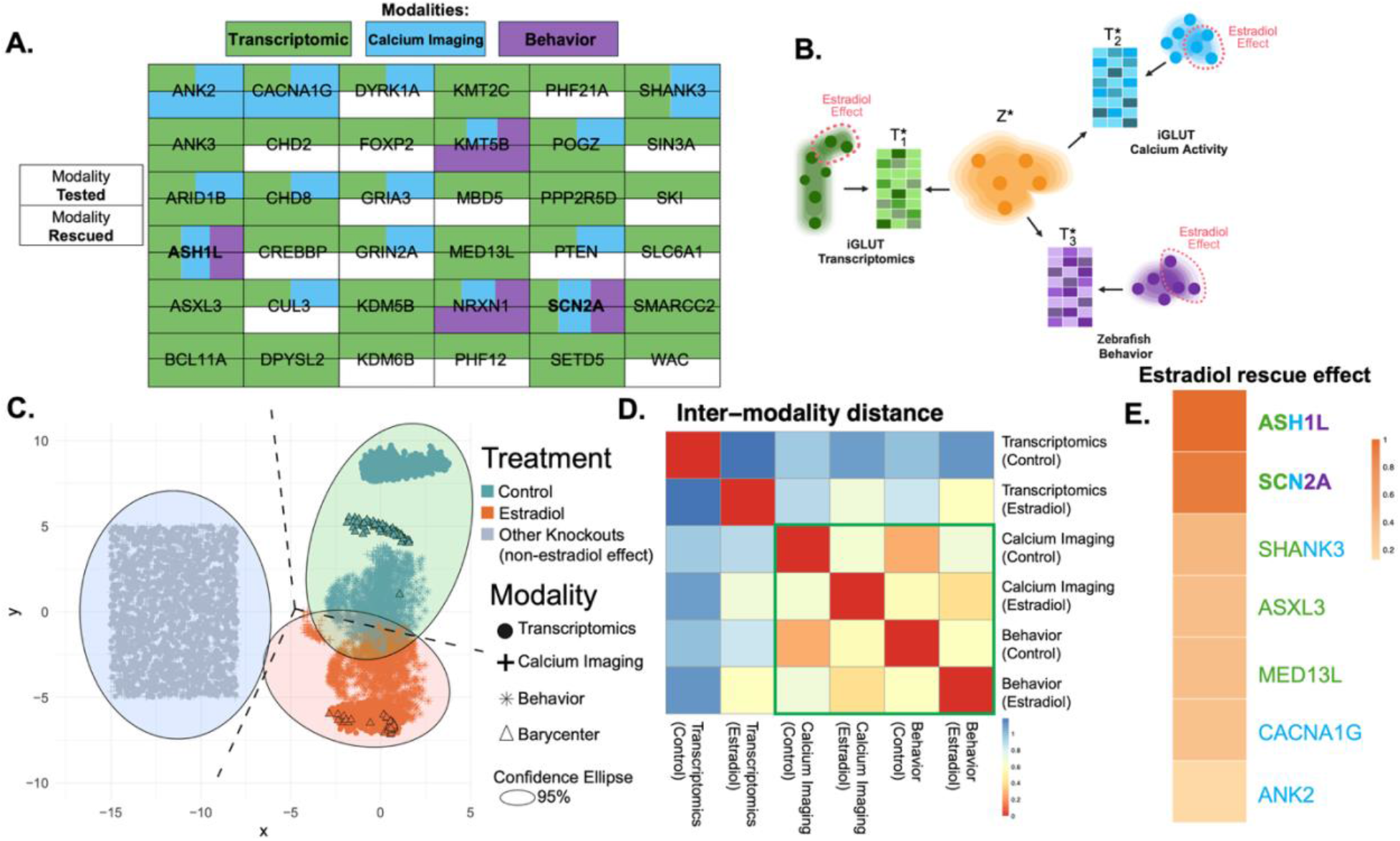
Multi-modal meta-analysis identifies greatest estradiol reversal effect in *ASH1L* and *SCN2A* mutants. **A.** Table of 36 NDD risk genes targeted across three modalities: iGLUT single-cell transcriptomics (green), iGLUT calcium imaging (blue), and zebrafish behavior (purple). For each target gene, the color(s) of the top half of the box indicates which modality was tested and color(s) of the bottom half indicates which modality estradiol rescued. White boxes indicate no estradiol rescue was observed in a mutant. **B.** Schematic of the Wasserstein Barycenter method of determining the geometric mean of each modality and separation of the estradiol effect from the control effect. T* indicates alignment plan for mapping one distribution to another and Z* indicates the barycenter latent space where the distributions collectively meet. **C.** Barycenter plot showing the control effect (green), estradiol effect (orange), and non-estradiol effect (gray). Modalities are represented by different shapes. Circle (•) indicates differentially expressed genes per mutant. Plus sign (+) indicates synchronization pattern per mutant. Asterisk (*) indicates individual larvae behavioral responses. Barycenters are indicated with triangles (Δ). **D.** Heatmap of the distance between modalities in projected latent space. Warm colors indicate closer distance, and cooler colors indicate farther distance. Scale bar is normalized to each modality’s distance in projected space. **E.** Ranking of genes based on magnitude of estradiol reversal effect across modalities.

By ranking ASD/NDD gene KOs on the degree of estradiol reversal across modalities, *ASH1L* and *SCN2A* were most responsive, followed by *SHANK3* (**Fig. 6E**). More modest and modality specific rescues were observed for *ASXL3*, *MED13L, CACNA1G,* and *ANK2* (**Fig. 6E; SI Table 22**). Overall, we identified a subset of 7 ASD/NDD genes (*ASH1L, SCN2A, SHANK3, ASXL3, MED13L, CACNA1G, ANK2)* that were particularly responsive to estradiol treatment.

To assess whether estradiol responsiveness predicted sex-biased clinical effects within ASD, we analyzed the sex distribution of *de novo* protein-truncating variant (PTV)^76^ carriers analyzed separately in the most (n = 7) and the least estradiol-responsive (n = 29) ASD/NDD genes and found they did not differ (Fisher’s exact test, OR = 0.93, 95% CI = 0.70–1.25, *P* = 0.669, **SI Fig. 15A-B, SI Tables 23-24**). Likewise, clinical phenotypes across six standardized ASD/NDD phenotypes in male and female participants in SPARK carrying variants in the seven most-responsive versus the 29 least-responsive ASD/NDD genes show no significant sex-interaction by PTV sensitivity analysis (**SI Table 25; SI Fig. 15C**). Overall, we report no evidence that estradiol responsiveness drives sex-biased clinical effects.

We next considered the brain expression of the most (n = 7) and least (n = 29) estradiol-responsive genes. First, we compared baseline sex-biased post-mortem brain cortical expression (absolute male–female log2 fold change) in two post-mortem datasets^77^, finding no significant difference in either (UCLA, *P* = 0.333; BrainVar, *P* = 0.375; meta-analysis, *P* = 0.757) (**Fig. 7A, SI Table 30**). Although we likewise resolved no significant differences in developmental cortical expression trajectories (most responsive Spearman ρ = 0.29, *P* = 0.535; least responsive Spearman ρ = −0.75, *P* = 0.052) (**Fig. 7B**), one subtype of excitatory neurons (upper-layer-enriched maturing excitatory neurons, ExM-U) showed a notable selective cell-type developmental expression signature^78^ associated with the most estradiol-responsive genes (median z-score 1.59 versus −0.11; IQR, 0.956–1.87 versus −0.520–0.276; Wilcoxon rank-sum test, *W* = 189, *P* = 5.06 × 10^−4^, FDR = 0.0081) (**Fig. 7C, SI Tables 28-29**).

**Figure 7:**
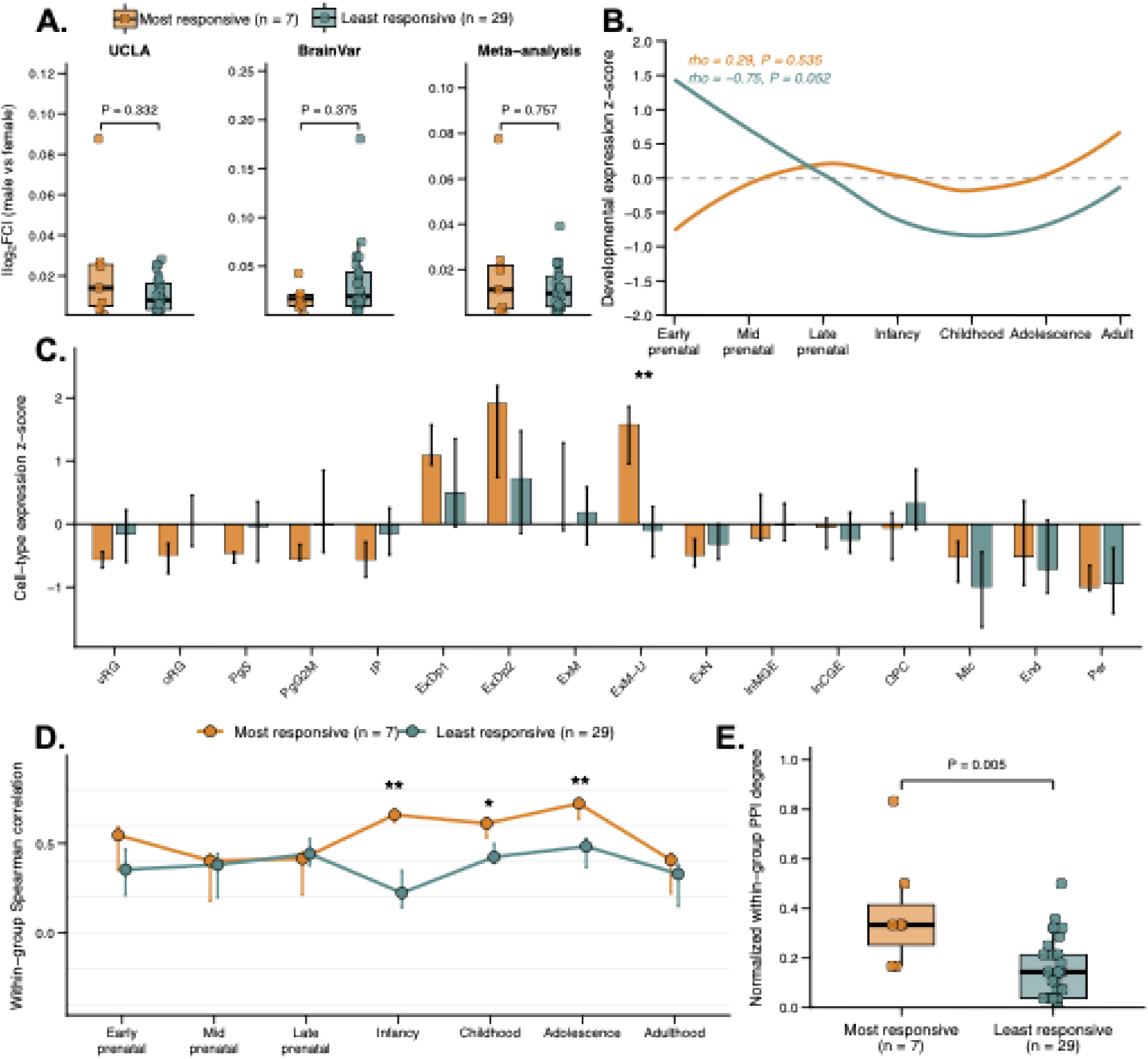
Network connectivity, developmental expression and sex-biased expression of estradiol-responsive genes. **A.** Magnitude of sex-biased gene expression in the UCLA and BrainVar cortical transcriptomic datasets and their meta-analysis. Absolute log2 fold change (|log2FC|) between male and female samples was used to quantify the magnitude of sex-biased expression. Boxes show the median and IQR, with individual points representing genes. Differences between the two gene groups were evaluated using two-sided Wilcoxon rank-sum tests. **B.** Developmental expression trajectories of the two gene groups in the BrainSpan cerebral cortex. Expression values were standardized within genes to z-scores, and curves show the developmental trajectories of the median gene-wise expression z-score. Spearman’s rank correlation coefficient (rho) and corresponding *P* values quantify the monotonic association between developmental stage and expression for each gene group. **C.** Cell-type-specific expression of the two gene groups in the developing human cortex. Gene-wise expression values were standardized across cell types, and bars represent the median cell-type expression z-score with interquartile range (IQR). Differences between the two gene groups were assessed using two-sided Wilcoxon rank-sum tests followed by Benjamini–Hochberg FDR correction across cell types. Cell-type abbreviations are as follows: vRG, ventricular radial glia; oRG, outer radial glia; PgS, S-phase cycling progenitors; PgG2M, G2/M-phase cycling progenitors; IP, intermediate progenitors; ExN, newborn/migrating excitatory neurons; ExM, maturing excitatory neurons; ExM-U, upper-layer-enriched maturing excitatory neurons; ExDp1 and ExDp2, deep-layer excitatory neuron subtypes 1 and 2; InMGE, interneurons derived from the medial ganglionic eminence; InCGE, interneurons derived from the caudal ganglionic eminence; OPC, oligodendrocyte precursor cells; Mic, microglia; End, endothelial cells; and Per, pericytes. **D.** Developmental-stage-specific co-expression connectivity of the two gene groups in BrainSpan cerebral cortex samples. For each developmental stage, pairwise gene–gene Spearman correlations were calculated across cortical samples, and the mean within-group Spearman correlation was calculated for each gene. Points indicate the median gene-level connectivity and error bars indicate the interquartile range (IQR). Group differences at each developmental stage were assessed using two-sided Wilcoxon rank-sum tests, with Benjamini–Hochberg false discovery rate (FDR) correction across stages. **E.** Comparison of normalized within-group protein–protein interaction (PPI) degree between the seven most estradiol-responsive genes and the 29 least-responsive genes using the STRING functional interaction network. Normalized degree was calculated as the number of within-group interactions for each gene divided by the maximum possible number of interactions within its gene group. Differences between groups were assessed using a two-sided Wilcoxon rank-sum test.

The most estradiol-responsive genes showed enhanced co-expression connectivity during postnatal cortical development, particularly during infancy (median connectivity, 0.660 versus 0.224; *P* = 2.36 × 10^−4^, FDR = 0.00165), childhood (0.611 versus 0.424; *P* = 0.0118, FDR = 0.0275) and adolescence (0.723 versus 0.484; *P* = 0.00159, FDR = 0.00557) (**Fig. 7D, SI Tables 26-27**). The most estradiol-responsive genes also showed significantly greater normalized within-group PPI connectivity than the least-responsive genes (mean normalized degree, 0.381 versus 0.160; Wilcoxon rank-sum test, *W* = 172.5, *P* = 0.0046, **Fig. 7E**), indicating stronger functional connectivity among the most-responsive genes. Together, these analyses indicate that, rather than sex-biased cortical expression, the seven most estradiol-responsive ASD/NDD genes showed enhanced co-expression during postnatal developmental periods and stronger functional network connectivity.

## DISCUSSION

Our multi-modal approach leveraged *in vitro* human cellular and *in vivo* zebrafish mutants to interrogate the interaction of estradiol with LOF mutations in functionally distinct and highly penetrant ASD/NDD risk genes. Overall, our data suggest that estradiol has both convergent and divergent effects across ASD/NDD risk genes, which are likely mediated by both genomic and non-genomic mechanisms. Consistent with earlier reports demonstrating estradiol suppression of ASD/NDD gene-associated phenotypes^31–34^, we report that estradiol ameliorates phenotypes associated with ASD/NDD LOF for select genes and identify *ASH1L* and *SCN2A* as most responsive to estradiol rescue. At the same time, our results underscore the importance of stratification of ASD genes^79,80^, given that some show substantially greater amenability to estradiol rescue across contexts. Overall, our multi-modal approach allows us to group ASD/NDD genes based on shared transcriptomic, circuit, and behavioral responses to estradiol across modalities.

Our analyses reveal broad transcriptomic convergence across ASD/NDD genes following estradiol exposure. Estradiol broadly ameliorated gene expression patterns resulting from all KOs examined (**Fig. 2**), converging on dysregulated neuronal gene and pathway expression independent of genotype, rescuing genes linked to neurotransmission, neurodevelopment, and cytoskeleton regulation, aligning with established biology^4,5,8–11,72,81,82^ (**SI Tables 31-32**). Similarly, estradiol rescued neurogenesis, synaptic, cell cycle, and metabolic, as well as estrogenic pathways in zebrafish *scn1lab* mutant brains. A plausible interpretation is that genomic mechanisms, at least in part, up-regulate genes that are functionally similar to ASD/NDD genes^83^ or their convergent downstream functions^43^. However, rescued genes were not more enriched for ER target genes, indicating that many effects downstream of estradiol likely also involve non-genomic mechanisms. Overall, our results support a model in which estradiol exposure modulates dysregulated gene expression in pathways that have been broadly implicated ASD/NDD, including neurogenesis, synaptic biology, and metabolism^25,35,43,59,84^, with stronger rescue effects in some genes compared to others, and with *ASH1L* and *SCN2A* showing the most pronounced rescue.

Strikingly, circuit- and behavioral-level analyses revealed genotype-specific reversals, and given their rapid timeframe, implicate non-genomic signaling. Estradiol broadly ameliorated gene expression patterns resulting from all KOs examined (**Fig. 2**), while selectively dampening network hyperexcitability across five genes: *ASH1L*, *SCN2A, ANK2, CACNA1G, SHANK3* in iGLUTs (**Fig. 3**). These five genes were all previously associated with seizure disorders in humans^85–91^ and LOF mutations in three resulted in hyperexcitability in glutamatergic neurons (*ASH1L*^92^*, SCN2A*^63^*, ANK2*^87^). Based on the immediacy of estradiol rescue of these select genes *in vitro*, here our data align with the observations of rapid non-genomic effects of estradiol on intracellular signaling and calcium mechanisms^19^. These mechanisms implicate calcium and MAPK signaling pathways but are less clearly linked to ASD/NDD^93^.

Our zebrafish studies support the stratification of ASD/NDD genes, demonstrating clustering of mutant behavioral phenotypes into subgroups with predicted differential responses to estradiol. Estradiol suppressed sleep-wake phenotypes in *ash1l* and *scn1lab* mutants (**Fig. 4**), consistent with estrogen rescue effects on nighttime hyperactivity in *cntnap2* mutants^31^. Estradiol also rescued regional baseline brain activity differences in a glutamatergic neuron cluster in the preoptic area in *scn1lab* mutants (**Fig. 5**). Interestingly, estradiol regulates sleep-wake patterns through the median preoptic nucleus via ERα and ERβ signaling in a sex-specific manner in rodents^94^. As a next step, live whole-brain calcium imaging during estradiol exposure could yield a more direct *in vivo* readout of its effects. Estradiol likewise suppressed induced seizures in *scn1lab* mutants (**Fig. 5**), which aligns with studies showing anti-seizure effects of estrogens^15,95^, although estradiol has seizure-promoting effects in other contexts^96^. Therefore, elucidating the interactions between genotype and estradiol on sleep and seizures in the context of select ASD genes is an important avenue for future research.

The seven most estradiol-responsive ASD/NDD genes do not predict sex-biases in clinical presentation or cortical brain expression (**Figs. 6-7**). We find no evidence that the prominent 4:1 male sex bias in ASD^97^ may be attributed to sex hormone levels. Indeed, circulating blood estrogen^98,99^ and brain estradiol concentrations are largely similar between males and females^100,101^; if anything, clinical evidence suggests that increased fetal steroidogenic activity correlates with ASD, rather than protecting against it^30,102,103^. The female protective effect model^104^ proposes that females require an increased genetic load of high-impact de novo variants^26,79^ and polygenic risk^105^ to reach the threshold for ASD, but this model is imperfect^106^ and confounded by ascertainment bias^107^. As new evidence reveals sex-differential trajectories in ASD^108^, accounting for later diagnosis of ASD in females markedly attenuates observed sex bias^109^. Beyond a handful of risk genes with evidence of sex-skewed association^42,79,110^, individual risk genes and polygenic risk scores confer similar liability for ASD in males and females^25,111^, particularly so for genes associated with severe developmental disorders, where male bias falls to 1.4:1^112^.

Although estradiol receptors are expressed at relatively modest levels, and only in a subset of human neurons (**SI Fig. 2)**^113,114^, neuronal estradiol levels likely exceed those in blood^115,116^. Estradiol is synthesized in the brain and localized to presynaptic terminals^117^, which may facilitate diverse mechanisms of estradiol signaling not observed in other tissues; for example, membrane bound ER *α* and ER *β* activate metabotropic glutamate receptors (mGluRs) independently of glutamate release, resulting in rapid ERK and CREB phosphorylation^72,118,119^. Our results indicate that complex interactions occur between ASD risk genes and estradiol responsive pathways in the brain (reviewed^120^). The enhanced co-expression and stronger functional network connectivity of estradiol-responsive ASD/NDD genes (**Figs. 6-7**) suggest points of downstream convergence that might be therapeutically actionable^43^. Novel precision therapeutic targets downstream of non-genomic estradiol receptor signaling may benefit a subset of individuals and warrant population-level investigations of the extent to which hormonal changes across the lifespan interact with genotype to impact traits linked to hyperexcitability, such as sensory sensitivity and sleep disturbances.

Precision psychiatry seeks to tailor treatments to individual patients, applying patient genotype, brain imaging, or cellular response to select targeted treatments^121^. Towards this, here we applied a multi-modal approach to stratify ASD/NDD risk genes based on shared transcriptomic, circuit, and behavioral response to estradiol. Our overarching hypothesis was that cellular and behavioral responses might inform clinical outcomes. Our study serves as a model to understand mechanisms of estradiol rescue of impacts associated with large-effect LoF variants in ASD/NDD risk genes. Although we focus on characterizing convergent and divergent estradiol responses in LoF variants, future work may extend to examining estradiol effects on common variants and incorporating pathway-based polygenic risk scores. An outstanding question, beyond the scope of this present study, is whether estradiol response is specific to a subset of ASD/NDD genes, or more likely, if any trait enriched for synaptic localized proteins would also have a subset of ER genes. Would any large set of genetic perturbations of brain expressed genes uncover estradiol response? Or is this specific to NDD/ASD genes? We interrogated whether estradiol-based rescue was gene-specific or generalizable and found that not every genetic risk factor is equally susceptible to estradiol. Particularly, we observe both gene-dependent convergent rescue and divergent exacerbation of behavioral phenotypes. Of note, estradiol rescue was not predicted by functional classification, as the top two genes had no overlap in annotated roles: *ASH1L* (histone methyltransferase) and *SCN2A* (sodium ion channel). Rather, estradiol acted to rescue shared downstream molecular, circuit, and behavioral phenotypes. Moreover, genes that are most susceptible to estradiol rescue show stronger functional network functional connectivity and are co- expressed at specific postnatal developmental windows. Together with our finding that these estradiol responsive genes are specifically enriched in maturing upper layer excitatory neurons, our study suggests that the ability of estradiol to rescue circuit- and behavioral-level phenotypes might represent a clinically actionable therapeutic window.

## METHODS

### Human stem cell models

#### Gene selection and CRISPR knockout library design

From the 102^25^ ASD/NDD, 72 ASD^26^, 309 DD^26^ and syndromic ASD^122^ datasets, we selected high-confidence NDD risk genes, defined as SFARI Gene (syndromic or score 1-2 genes)^122^ or classified as “definitive” in the Gene2Phenotype (G2P) database^123^. Gene selection prioritized loci with elevated expression during early human cortical development based on single-nucleus RNA-seq data ^124^data and/or BrainSpan^125^. For arrayed CRISPR screening (calcium imaging experiments), we targeted 15 genes spanning major functional categories: chromatin remodeling & transcriptional regulation (*ASH1L, POGZ, ARID1B, KMT5B, CHD8*), synaptic function (*NRXN1, SHANK3, CACNA1G, GRIA3, GRIN2A, SCN2A*), cell signaling & protein modification (*DYRK1A, PTEN, CUL3*), and cytoskeletal proteins (*ANK2*). For pooled CRISPR screening (estradiol treatment and scRNA-seq experiments), we expanded to 36 genes: chromatin remodeling & transcriptional regulation (*ASH1L, POGZ, ARID1B, KMT5B, CHD8, CHD2, SMARCC2, BCL11A, SKI, ASXL3, CREBBP, FOXP2, KDM5B, KDM6B, KMT2C, MBD5, MED13L, PHF12, PHF21A, SETD5, SIN3A, WAC*), synaptic function (*NRXN1, SHANK3, CACNA1G, GRIA3, GRIN2A, SCN2A, SLC6A1*), cell signaling & protein modification (DYRK1A, PTEN, CUL3, PPP2R5D), and cytoskeletal proteins (*ANK2, ANK3, DPYSL2*). Gene selection was not based on sex-differential penetrance; a subset of genes (e.g. *MBD5, ASH1L,* and *CUL3)* are male-biased, others show enrichment in both males and females^42,110^ (**SI Table 1**).

Each gene was targeted with 3-4 gRNAs (LentiArray Custom CRISPR Library, ThermoFisher, #A26620) or scrambled controls (LentiArray CRISPR Control Lentivirus Particles, ThermoFisher, #A32062). Lentiviral libraries were generated by ThermoFisher. For arrayed screening, 250,000 hESC-iCas9 cells per well were transduced overnight at MOI >1. For pooled screening, cells were transduced at MOI <0.2 to ensure single integration events. Following transduction, cells were selected with 1 µg/mL puromycin for 48 hours until control wells were cleared, then expanded in mTeSR Plus and cryopreserved.

#### NPC culture

Validated control hiPSC-derived NPCs were selected from a previously reported case/control hiPSC cohort of childhood onset SCZ (COS)^44^: NSB2607-2-1 (karyotypic XY, average SCZ PRS, European ancestry), NSB3182-3-1 (karyotypic XX, average SCZ PRS, European ancestry). hiPSC-NPCs were generated via dual-SMAD inhibition (0.1µM LDN193189 and 10µM SB542431) followed by neural rosette selection and MACS-based purification and validated as previously described^44^ and cultured in hNPC media (DMEM/F12 (Life Technologies #10565), 1x N2 (Life Technologies #17502-048), 1x B27-RA (Life Technologies #12587-010), 1x Antibiotic-Antimycotic, 20ng/ml FGF2 (Life Technologies)) on Matrigel (Corning, #354230).

For pooled CRISPR screens, hiPSC-NPCs were transduced with lentiviral-Cas9v2 (Addgene #98291) and LV-gRNA (ThermoFisher #A31949) and 24-hours later selected with 1 mg/mL puromycin (Sigma, no. P7255) and 1mg/mL hygromycin as described previously^44^. 72-hours prior to harvest, NPC were transduced with LV-gRNA, 24 hours prior to harvest, NPCs were treated with estradiol.

#### iGLUT neuron induction

The H1 (NIHhESC-10-0043, XY) inducible Cas9 (iCas9)^46^ cell line was used to generate NGN2-induced iGLUT neurons for pooled Perturb-seq of 36 NDD-associated genes and 15 individual knockout lines for calcium imaging. gRNAs were delivered to H1 iCas9 hESCs prior to differentiation (pooled or individually), with Cas9 activation and knockout occurring during doxycycline-inducible NGN2 conversion to iGLUT neurons.

Undifferentiated hESCs were maintained in mTeSR Plus media (#100-0276, STEMCELL Technologies) with complete media changes performed daily. At 85-90% confluency, cells were either passaged using Accutase (#AT-104, Innovative Cell Technologies) for single-cell passaging and plated at 15,000 cells/cm² in 6-well plates coated with Geltrex (#A1413302, Gibco), or dissociated with 0.5 mM EDTA and split every 3-4 days at a 1:10 ratio. hESCs were maintained at 37°C with 5% CO₂. Mycoplasma contamination was routinely monitored using the MycoAlert PLUS Mycoplasma Detection Kit (#LT07-703, Lonza), and all cell lines tested negative.

H1-iCas9 hESCs^46^ were transduced with a lentivirus expressing doxycycline-inducible *NGN2* (pLVX-UbC-rtTA-Ngn2:2A, Addgene #127288) modified to express blasticidin resistance in place of EGFP. Cells were plated at 100,000 cells/cm² and selected with 10 µg/mL blasticidin for 5 days. Two million selected cells were seeded per 10 cm dish (35,000 cells/cm²) and cultured in neural induction medium (NIM; **SI Table 19**). NGN2 expression was induced with 2 µg/mL doxycycline for 72 hours (day 0-3) with continued blasticidin selection until day 4. On day 4, cells were dissociated and replated onto Geltrex-coated 12-well plates at 100,000 cells/cm² in neural maturation medium (NMM; **SI Table 19**) supplemented with 10 µM Y-27632 ROCK inhibitor. From day 5 onward, neurons were maintained in NMM with half-medium changes every 2-3 days.

#### 17β-estradiol treatment

During fetal corticogenesis, clinical reference levels for circulating maternal estradiol levels range from 1,561 pg/mL (5.7 nM) to >30,000 pg/mL (110 nM)^126^, of which approximately one-third is considered to be bioavailable (free or albumin-bound)^98^. Cortical estradiol levels generally exceed those found in circulating blood^115,116^, with measurements in adult males reported to be nearly 10 nM^127^. Studies of rapid, nongenomic effects in mouse, rat, primate, and human neurons in the literature indicated that increasing estradiol treatment from 100pM to 100 nM increased the fraction of responding neurons^128^. Treatment of neurons with 1-100 nM estradiol rapidly altered actin polymerization, vesicle release probability, spine density, and synaptic plasticity, with effects that vary by neuron identity, receptor expression, and sex^8–10,12–14,129^. NPCs or iGLUT neurons were treated with 17β-estradiol (E2; Sigma-Aldrich, #E2758) at 100 nM or vehicle control (0.01% DMSO), acutely for calcium imaging or for 24 hours prior to single-cell capture.

##### Definition of exacerbated, unchanged, partial rescue, rescue, and over-corrected

In NPCs and iGLUTs, for DEGs and cellular phenotypes that are significantly different in the KO+DMSO vs non-targeting control (NT)+DMSO (“KO-WT”) p-value ≤ 0.05, we characterize the effect of the KO-x-E2 on a feature. If KO+E2 vs NT+DMSO (“KO-E2”) p-value >0.05, the feature is classified as *rescued.* If “KO-E2” p-value ≤ 0.05, we evaluate KO+E2 vs KO+DMSO (“E2-Effect”). If the “E2-Effect” p-value > 0.05, the feature is classified as *unchanged*. If the “E2-Effect” ≤ 0.05, we evaluate scaled beta values of “KO- WT” and “KO-E2”. If scaled beta values are in opposite directions, the feature is classified as *over-corrected*. If scaled beta values are in the same direction, we evaluate the absolute values of the beta scores. If the absolute value of the scaled beta of “KO-E2” > absolute values of scaled beta of “KO-WT”, the feature is classified as *exacerbated*. If the absolute value of the scaled beta of “KO-E2” < absolute values of scaled beta of “KO- WT”, the feature is classified as *partially rescued*. The degree of rescue thresholds decision tree is illustrated in **SI Fig. 16** and reproduced here for clarity.

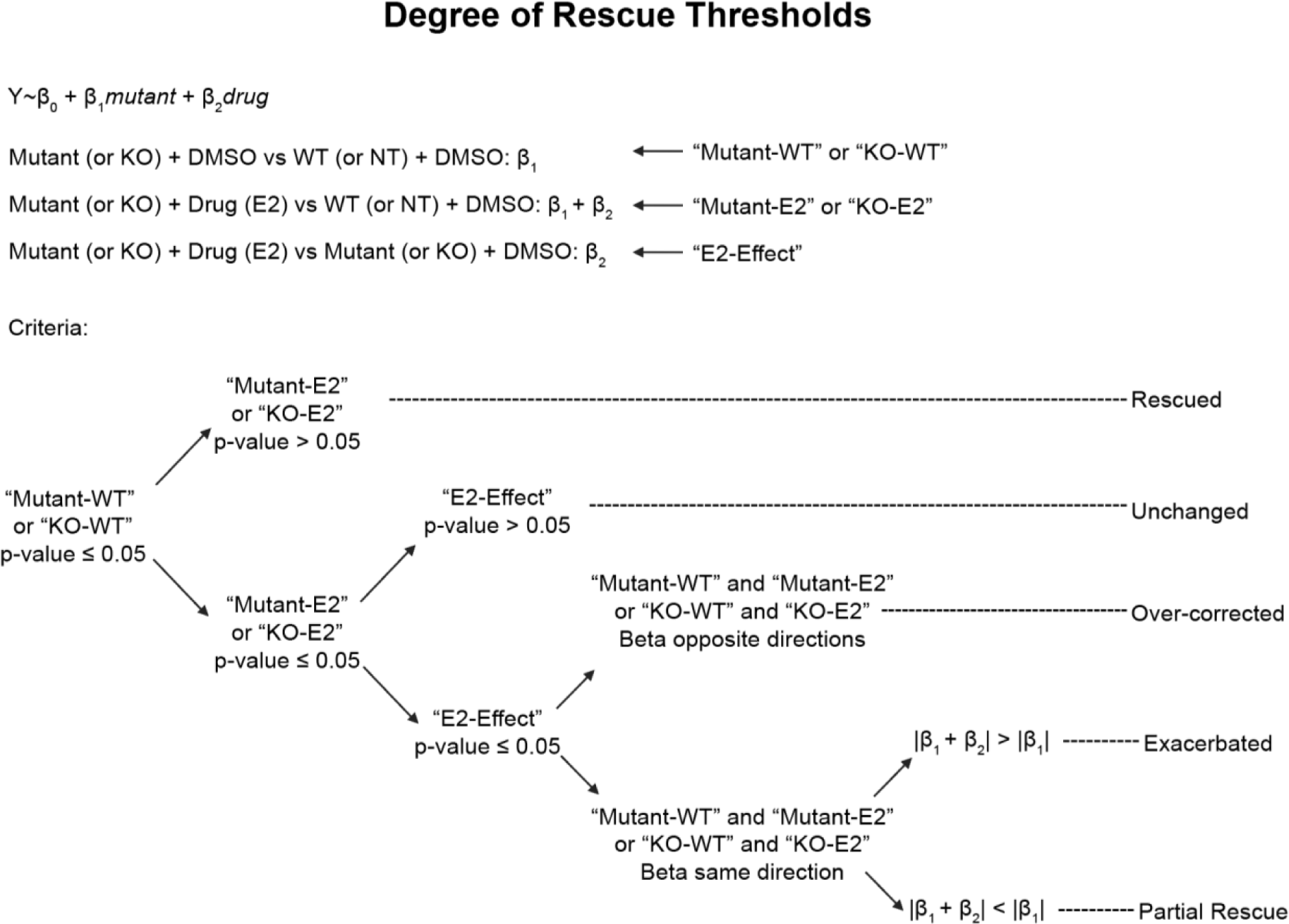

#### Single-cell RNA sequencing library preparation

After 24 hours of estradiol/vehicle treatment, NPCs or iGLUT (22DIV) cultures were washed once with DMEM and dissociated with Accumax (Innovative Cell Technologies) at 37°C with gentle trituration. Cell suspensions were filtered through 37 µm cell strainers, counted, and pelleted at 500 × g for 5 minutes. Cells were resuspended in eBioscience Flow Cytometry Staining Buffer (#00-4222-26, Invitrogen) and adjusted to 2.2 × 10⁶ cells/mL with viability >80% as determined by trypan blue exclusion. A targeting recovery aimed at ∼52,000 cells per lane (83,000 cells loaded). Single-cell capture and library preparation were performed using the Chromium Single Cell 5’ v3 workflow with Feature Barcode technology for CRISPR Detection (10x Genomics) according to manufacturer’s protocol at Keck Microarray Shared Resource (KMSR) at Yale University. cDNA was amplified for 10 PCR cycles, and CRISPR guide capture libraries were generated using 10 PCR cycles for the guide RNA enrichment product.

#### Sequencing and data processing

Library quality was assessed using a Fragment Analyzer. Sequencing was performed on a NovaSeq 6000 (Illumina) at Yale Center for Genome Analysis (YCGA). Gene expression (GEX) libraries were sequenced to a depth of ∼30,000 reads per cell, and CRISPR guide libraries to ∼5,000 reads per cell.

#### Analysis of single-cell CRISPRko screens

Analysis of single-cell transcriptomics data was performed using the Seurat V5^130^ R package, with quality control procedures conducted independently for each cell type and treatment dose. In the initial QC phase, percentage of mitochondrial, ribosomal, and hemoglobin genes were calculated and filtered based on cell type-specific thresholds.

Further filtering was applied based on RNA counts, detected features, gRNA counts, and detected gRNA features. To assign guide RNAs, gRNA counts were normalized via the Seurat::NormalizeData function and then demultiplexed using Seurat::MULTIseqDemux. Only cells identified as singlets with a single assigned guide were retained for downstream analyses. For NPCs specifically, an additional step to reassign certain doublets as singlets was performed to maximize the number of cells for downstream analyses: doublets that contained non-targeting and a KO guide were assigned the KO guide, and all other doublets that were not singlets were removed. Following the identification of singlets, an additional round of cell and gene filtering was performed: cells were excluded based on UMI count, number of genes detected, and log10GenesPerUMI. Ribosomal, mitochondrial, hemoglobin, and MALAT1 genes were also removed from the dataset. To account for cell-type specificity and maturity, cell-type marker scores were calculated using the Seurat::AddModuleScore function and visually validated across clusters. Normalization and regression of biological covariates—including mitochondrial & ribosomal & hemoglobin gene percentages, cell cycle phase, subtype scores (iGLUT), donor (NPC), and astrocyte & neural crest cell scores (NPC)—were performed using Seurat::SCTransform. From the remaining cell population, gRNA-specific differential expression analysis was performed against cells containing scrambled gRNAs. Only guides that produced significant downregulation of the target gene (p < 0.1 & logFC < 0) were retained. Cells were then collapsed by gene target identity for WNN, and the resulting clusters were designated as dominant or heterogeneous clusters using the following criteria. Clusters in which a single gRNA represented >75% of cells were assigned to that gRNA. All other clusters were initially classified as Mixed. When the Scrambled control gRNA was present, it was treated separately: clusters in which Scrambled represented >50% of cells were classified as Mixed. Otherwise, Scrambled cells were excluded, gRNA proportions were recalculated, and the cluster was reassigned to a single gRNA only if one remaining gRNA exceeded the 75% threshold; if not the cluster remained classified as Mixed. All cells that did not fall into a single gRNA cluster were removed from downstream analyses. Gene targets were included in the final analysis only if more than 75 cells remained across all treatment conditions. Prior to downstream analyses, genes on the Y chromosome were removed, and integration of the treatment conditions were performed using Seurat::IntegrateData. Differential expression analysis between conditions were performed using Seurat::FindMarkers. All gene ontology enrichment were done using clusterProfiler^131^::gseGO.

#### Meta analysis of perturbation effects on gene expression

Summary statistics from differential expression analysis between KO and Non-targeting cells across the 16 KOs were supplied to METAL^132^. “Convergent” genes were defined as those with significant meta-analysis FDR (FDR < 0.05), non-significant Cochran’s heterogeneity Q test p (p > 0.05), and shared direction across all 16 KOs^43^. “Nonconvergent” genes were defined as those that did not meet the convergence threshold.

#### Enrichment test for ER targets and E2 amelioration

Fisher’s exact test was used to test the enrichment of ER targets or E2 rescue genes in convergent genes. Estrogen receptor targets were pulled from multiple sources (Ingenuity Pathway Analysis^133^ database: IPA_ESR1, IPA_ESR2; ENCODE Transcription Factor Targets^134,135^: ENCODE_ESR1; ChEA^136^ Transcription Factor Targets: CHEA_ESR2; LINKS L1000 CMAP CRISPR Knockout Consensus Signatures^137^: GPER1). IPA gene sets were exported from the IPA software pathway build module, and all other datasets were downloaded from Harmonizome 3.0^138,139^.

#### Calcium Imaging

The calcium indicator, fluo4-AM was introduced into neuronal networks at TD30 with 2μM. All imaging was performed immediately after incubating with fluo4-AM for 1hr. Calcium transients were recorded at an excitation of 488nm using a x20 water-immersion objective (Leica) with 1024 x 1024- pixel resolution and 0.6 x 0.6- mm field of view at rate of 80ms/frame by a Dragonfly spinning disc microscope. Recording was performed in artificial cerebrospinal fluid (aCSF): NaCl: 130mM; KCl: 3mM; NaH2PO4: 1.25mM; NaHCO3: 26mM; MgCl2: 1mM; CaCl2: 1mM with additional D-(+)-glucose: 5mM. All samples were subjected to incubation with E2 (Sigma-Aldrich, #E2758) at 100nM and the identical fields were reimaged after E2 exposure.

Raster plots were generated for all the identified cells/ regions of interests in the calcium imaging video. The orange dots represent calcium activity or spike in a specific cell and a specific time frame. Each block along the y axis represents a cell, and each block along x axis represents a time frame. All raster plots were generated in MATLAB and then subjected to hierarchical clustering using UPGMA, via hclust function in R. The heatmaps were generated using the ‘pheatmap’ package in R.

The calcium imaging analysis is performed by taking raw calcium fluorescence intensity for every pixel across all time frames. After this step, the dominant pixels are chosen after dimensionality reduction approach (via Principal Component Analysis), where the pixels showing the most variation with time are filtered. After this, the temporal noisiness is removed deconvolving persistent and lagging calcium signals to get sharp peaks. After removing noise, the dominant pixels are also filtered spatially for each time frame to keep the pixels showing the most variation. After this, spiking cells are identified by looking at changes in intensity patterns in the spatial neighborhood of the selected pixels. The identified cells are further checked for overlapping calcium signals by spatial deconvolution, giving rise to unique cells. After the cells are identified, a machine learning algorithm (Markov-chain Monte-Carlo) is used to train the basal calcium signal over time. Later, these basal signals are used to identify signals which fluctuate far beyond the basal level, termed as spikes. These calcium spikes are identified for every time point and later merged temporally to identify all time-continuous spikes. The calcium spikes are characterized for all the identified cells, and later, spike characteristics like, spike amplitude (height of the spike from basal level), spike frequency, spike duration, and spike counts (total spike per cell during the length of imaging) are calculated and recorded for further analysis^140,141^.

#### Multielectrode Array

Multi-electrode array (MEA) extracellular neuronal activity recordings were performed using the Axion Maestro system (Axion Biosystems) at a sampling frequency of 12.5 kHz. Signals were acquired in Neural Spikes mode, median referenced and filtered using a 200-Hz IIR high-pass filter and a 3-kHZ Kaiser-window low pass filter. Spikes were detected using the AxIS Adaptive Threshold Crossing algorithm with a variable detection threshold set at 5.5 standard deviations above the electrode specific noise level. Electrodes with a minimum firing rate of 5 spikes/min were considered active. Bursts were detected using an inter-spike-interval (ISI) threshold algorithm and were defined as a minimum of 5 spikes with a maximum ISI of 100ms. Network bursts were defined as a minimum of 50 spikes with a maximum ISI of 100ms and participation of at least 35% of electrodes. Synchrony was assessed using a 20-ms synchrony window. MEA measurements were quantified using Axion analysis software.

A total of 150,000 induced glutamatergic (iGLUT) neurons were seeded per MEA well, which had been double-coated with poly-D-lysine and Geltrex. Half-medium changes were performed every 2–3 days. MEA measurements were conducted following 24 hours of treatment with either DMSO (vehicle control) or estradiol (100nM) at the same final volume ratio, using D21 iGLUT cultures.

#### Zebrafish

All procedures involving zebrafish were conducted in accordance with Institutional Animal Care and Use Committee (IACUC; Protocol #2024-20054) regulatory standards at Yale University. Zebrafish larvae were raised at 28°C on a 14:10 hour light:dark cycle. Larvae were grown in 150 mm Petri dishes in blue water (0.3g/L Instant Ocean, 1 mg/L methylene blue, pH 7.0) at a density of 60-80 larvae per dish. Behavioral assays were conducted in zebrafish larvae at 5-7 dpf. At these developmental stages, sex is not yet determined.

#### Zebrafish mutant generation

Stable zebrafish lines were generated by our lab (*arid1b^Δ7/Δ7^*, *chd8^Δ7/Δ7^, kdm5ba^Δ17/Δ17^b^Δ14/Δ14^*, *pogza^Δ23/Δ23^b^Δ20/Δ20^, cul3a^Δ7/Δ7^b^Δ32/Δ32^, dyrk1aa^Δ77/Δ77^b^Δ8/Δ8^, scn1lab^Δ44/Δ44^, nrxn1a^5i/5i^b^10i/10i^, ptena^Δ10/Δ10^b^Δ2/Δ2^*)^38,43,59^, or provided as a generous gift from the Thyme lab (*ash1l^1i,Δ60,19i/1i,Δ60,19i^, kmt5b^Δ208,1i,Δ5/Δ208,1i,Δ5^, kmt2ca^Δ82,17i/Δ82,17i^b^Δ6,Δ29/Δ6,Δ29^, grin2aa^Δ175Δ175^,grin2ab^Δ42Δ5Δ38/Δ42Δ5Δ38^, cacna1g ^Δ52/Δ52^)*^53,55,142^ and the Dallman lab (*shank3a^4i^shank3b^1i^)*^54^. F0 crispants (*ank2ab, bcl11aab, chd2, crebbpab, gria3ab, kdm6bab, mbd5, phf12ab, phf21aab, ppp2r5d, setd5, sin3aab, skiab, slc6a1ab smarcc2, wacab)* were generated as^52^. Briefly, we designed two CRISPR crRNAs per allele, prioritizing early exons for targeting. CRISPR RNPs were assembled individually and then combined prior to injection at the one-cell stage. The number of scrambled guides injected into the control group was matched to the number of CRISPR guides used for the experimental group. Injected embryos were raised to 5 dpf at which point the behavioral assays (described below) were conducted. We identified unique behavioral fingerprints for each NDD gene mutant, revealing convergent and divergent phenotypes across mutants (**SI Fig. 5A**). To classify convergent behavioral subgroups that may share circuit-level functions, we performed correlation analyses with hierarchical clustering across mutants. We identified seven distinct subgroups of NDD genes with highly correlated behavioral features (**Fig. 4C**).

#### Behavioral assays

We performed automated, high-throughput, quantitative behavioral profiling of larval zebrafish to measure arousal and sensorimotor processing as a readout of circuit-level deficits resulting from gene perturbation.^38^ We quantified 24 visual-startle^38,56^ and sleep- wake parameters^57,58^ responses in 15 stable homozygous mutant or F0 mosaic crispant lines for 16 NDD genes (**SI Tables 8-9**). Larvae were placed into individual wells of a 96 well plate (7701-1651; Whatman, Clifton, NJ) containing 650 µL of standard embryo water (0.3 g/L Instant Ocean, 1 mg/L methylene blue, pH 7.0) per well within a Zebrabox (Viewpoint LifeSciences; Viewpoint Life Sciences, Montreal, Quebec, Canada). Locomotion was quantified with automated video-tracking system (Zebrabox and ZebraLab software). The visual-startle response (VSR) assay was conducted at 5 days post fertilization (dpf) as described^38^. To assess larval responses to lights-off stimuli (VSR-OFF), larvae were acclimated to white light for 1 hour, and baseline activity was tracked for 30 minutes followed by five 1-second dark flashes with intermittent white light for 29 seconds. To evaluate larval responsivity to lights-on stimuli (VSR-ON), the assay was reversed, where larvae were acclimated to darkness for 1 hour, and baseline activity was tracked for 30 minutes followed by five 1-second white light flashes with intermittent darkness for 29 seconds. For VSR-OFF and VSR-ON, six behavioral parameters were quantified using custom MATLAB code^38^ (available on github at https://github.com/ehoffmanlab/Weinschutz-Mendes-et-al-2023-behavior; DOI:10.5281/zenodo.7644898): (i) average intensity of all startle responses; (ii) average post-stimulus activity; (iii) average activity after first stimulus; (iv) stimulus versus post-stimulus activity; (v) intensity of responses to the first stimulus; (vi) intensity of responses to the final stimulus. The sleep-wake paradigm was conducted between 5-7 dpf, following the VSR-OFF and VSR-ON assays. During a 14h:10h white light:darkness cycle, larvae activity and sleep patterns were tracked within the Zebrabox and analyzed with custom MATLAB code^57^ (available at (https://github.com/JRihel/Sleep-Analysis/tree/Sleep-Analysis-Code; <u>DOI: 10.5281/zenodo.7644073</u>). Six behavioral parameters were quantified for daytime and nighttime: (i) total activity; (ii) total sleep; (iii) waking activity; (iv) rest bouts; (v) sleep length; (vi) sleep latency. Across VSR-OFF, VSR-ON, and sleep-wake assays, we analyzed 24 parameters.

#### Behavioral analysis

Linear mixed models (LMM) were used to compare phenotypes of each behavioral parameter between homozygous mutant versus wild-type or crispant versus scramble-injected fish for each gene of interest. Variations of behavioral phenotypes across experiments were accounted for by including the date of the experiment as a random effect in LMM. Hierarchical clustering analysis was performed to cluster mutants and behavioral parameters based on signed −log10-transformed p-values from LMM, where sign indicates direction of the difference in behavioral phenotype when comparing stable mutant to wild-type or crispant to scrambled-injected. Pearson correlation analysis was used to assess correlations between mutants based on the difference in the 24 parameters. Difference was evaluated using signed −log10-transformed p-values.

#### Drug prioritization based on zebrafish pharmaco-behavioral profiles

NDD gene-associated mutant and crispant behavioral phenotypes were compared to a dataset of 376 U.S. FDA-approved drugs that were screened for their behavioral effects in larval zebrafish using the visual-startle and sleep-wake assays described above ^59^. These drugs have a significant effect on at least two behavioral parameters (LMM, p<0.05/3, corrected for three behavioral assays). Pearson’s correlation analysis was used to identify drugs that significantly correlate (correlation >0.5, p<0.05, t-statistic) or anti-correlate (correlation <-0.5, p<0.05, t-statistic) with mutant behavioral signatures (**SI Table 14**).

#### Estradiol treatment in zebrafish

For mutant-x-estradiol experiments, larval activity was monitored from 5-7 dpf. For acute estradiol experiments, individual wild-type zebrafish larvae were added to each well of a 96-well plate containing 650 μl of standard embryo water. A 10 mM stock solution of *β*- Estradiol 17-cypionate (Millipore Sigma #E8004-1G) dissolved in DMSO or DMSO alone (control) was pipetted directly into each well at a final concentration of 20 µM (0.2% DMSO final concentration) in 12-24 background-matched homozygous or wild-type larvae after which the visual-startle and sleep-wake assays were performed. For chronic estradiol experiments, larvae were treated with 20 µM estradiol or DMSO in blue water in 150 mm Petri dishes for 24 h then pipetted into a 96 well plate of fresh 20 µM estradiol or DMSO water, after which visual-startle and sleep wake assays were performed. Genotyping was conducted after each experiment to confirm genotypes for stable mutant lines.

##### Definition of exacerbated, unchanged, partial rescue, rescue, and over-corrected (SI Fig. 16)

Z-test of drug rescue was used to assess statistical significance. For behavioral features with mutant p-value ≤ 0.05 (“Mutant-WT,” HOM DMSO vs WT DMSO), features with a drug rescue p-value >0.05 (“Mutant-E2,” HOM E2 vs WT DMSO) were considered *rescued*. Among the remaining features, those with a drug p-value >0.05 (“E2 Effect,” HOM E2 vs HOM DMSO) were considered unchanged. If the drug p-value <0.05, features in which the absolute drug rescue effect (“Mutant-E2,” HOM E2 vs WT DMSO) was in the opposite direction than the absolute mutant effect (“Mutant-WT,” HOM DMSO vs WT DMSO) were considered *over-corrected*. If the drug p-value <0.05, features in which the absolute drug rescue effect (“Mutant-E2,” HOM E2 vs WT DMSO) was in the same direction and smaller than the absolute mutant effect (“Mutant-WT,” HOM DMSO vs WT DMSO) were considered *partially rescued*, otherwise, they were considered *exacerbated*.

#### Immunohistochemistry

Whole mount antibody staining of dissected embryos was performed as described in ref. ^38^. To test the effect of compounds on baseline brain activity, *scn1lab^Δ44/Δ44^* and background-matched *scn1lab^+/+^* larvae were exposed to estradiol at a final concentration of 20 µM dissolved in DMSO or DMSO alone for 24 hours beginning at 5 dpf. Sample sizes: *scn1lab^+/+^* + DMSO (n=34), *scn1lab^Δ44/Δ44^* + DMSO (n=35), *scn1lab^Δ44/Δ44^* + estradiol (n=37) across 3 independent datasets. Larvae at 6 dpf were collected by pouring larvae into a mesh sieve and immediately immersing the sieve into 4% PFA/4% sucrose in PBS + 0.25% Triton (PBST). Larvae were fixed overnight at 4°C and washed in PBST. Brains were dissected prior to immunostaining, as in Wilson et al. 1990^143^. Dissected brains were incubated in 150 mM Tris-HCl for 15 minutes at 70°C, washed in PBST, permeabilized in 0.05% Trypsin-EDTA for 45 minutes on ice, and washed in PBST. Brains were blocked for one hour in PBST with 10% BSA, 2% normal goat serum (NGS) and 1% DMSO, and incubated overnight at 4°C in primary antibodies diluted in blocking solution without NGS. The following antibodies and dilutions were used: pERK (Cell Signaling #4370, RRID: AB_2315112) and tERK (Cell Signaling #4696, RRID:AB_390780), 1:500. Samples were then washed in PBST and incubated for 3 hours at room temperature with Alexa-fluorophore-conjugated secondary antibodies (Life Technologies) at a dilution of 1:200 in blocking solution without NGS. Immunostained brains were mounted dorsally in low melting point agarose (1-2%) and imaged by confocal microscopy at 25X (0.95W) using z-stacks ranging from ∼200- 300 μm (Stellaris 5, Leica Biosystems), with a step size of 1.5µm.

#### Whole brain activity mapping analysis

Whole-brain activity mapping was performed using a custom brain-mapping pipeline^144^ to quantify regional differences in brain activity. Custom software used in this analysis is available on github at https://github.com/ehoffmanlab/Jin_Neelakantan_BREEZE_Mapping; DOI: <u>10.5281/zenodo.7996403</u>). Briefly, confocal z-stack images of tERK- immunostained brains were nonlinearly registered to a standard reference brain^145^ using transformation-based deformation^146,147^. The geometric transformation is applied to the pERK confocal z-stack images. The pERK image intensities were normalized by tERK intensities through a voxel-wise division (pERK/tERK). The average pERK/tERK intensities were calculated within individual regions delineated in the Z-brain atlas^145^. The 4-region, 8-region and 149-region atlases were derived manually from versions of the Z-brain atlas^65,145^ as described in Weinschutz Mendes et al. (2023)^38^. Linear mixed modeling was applied to raw pERK/tERK values to identify regions in which mutant brain activity phenotypes were rescued following E2 exposure.

#### Visualization of brain activity phenotypes

Bioimage Suite Web^148^ was used to visualize differences in brain activity in estradiol or vehicle treated homozygous mutants versus wild-type background-matched, vehicle treated fish. Images for activity were overlayed on a resampled standard reference fish. We calculated a voxel-wise Z-score normalized value of mutant compared to background-matched wild-type larvae using the mean and standard deviation of pERK/tERK values. Images were generated in BioImage Suite Web to capture cross sectional increases and decreases in the axial and sagittal planes. Activity differences between drug- versus DMSO-treated samples were visualized as a subtraction of the normalized means. Activity differences were visualized by mapping Z-score values from 1-3 for *scn1lab*. All axial mosaics are shown starting at Z level of 80 and decreasing by 10 slices for each image. Sagittal mosaics are displayed starting at X level of 80 and decreasing by 10 slices for each image.

#### Brain activity analysis

LMM with the main effects of E2 and mutant were used to assess E2 and mutant effects on activity in each brain region using 3 sets of fish including DMSO-treated wild-type (*scn1lab^+/+^*, DMSO-treated), homozygous (*scn1lab^Δ44/Δ44^*) mutant, and E2-treated homozygous (*scn1lab^Δ44/Δ44^*) mutant. The variations of brain activity across experiments were accounted for by including the date of the experiment as a random effect in LMM. Drug rescue effects were defined as the combined effects of E2 and mutant on activity in each brain region to compare brain activities between E2-treated mutant and DMSO-treated wild-type fish. Z-test of E2 rescue was used to assess statistical significance. For regions with mutant p-value <0.05, regions with E2 rescue p-value >0.05 were considered rescued. Among the remaining regions, those with E2 p-value >0.05 were considered as no change. If E2 p-value <0.05, regions in which the absolute E2 rescue effect was smaller than the absolute mutant effect were considered as partial rescue, otherwise, they were considered as exacerbated. A visual of the degree of rescue thresholds decision tree is included in **SI Fig. 16**. We conducted independent LMM analysis of each dataset then pooled the estimation together across the 3 experiments to generate a pooled meta-analysis. The pooled estimate was calculated as the inverse-variance weighted average of the experiment-specific estimates. Statistical significance across experiments was assessed using Fisher’s combined probability test to obtain a pooled *p*-value. We classified the regions into different degrees of rescue based on the pooled estimates.

#### Seizures

PTZ-induced seizure assays were performed as described in Hoffman et al. (2016)^31^ with modifications. For estradiol rescue experiments, larvae were exposed to estradiol 20 µM or DMSO control, n=24 per genotype per group) over night from 4 dpf – 5 dpf in petri dishes. At 5 dpf, larvae were pipetted into estradiol (20µM) or vehicle water (650 L of standard methylene blue embryo water (0.3g/L Instant Ocean, 1 mg/L methylene blue, pH 7.0)) in the wells of a 96 well plate containing estradiol or vehicle water. The plate was placed in a custom-modified Zebrabox. Larval movement was recorded using Videotrack quantization mode for 30 minutes before and after the addition of PTZ or water. Videotrack detection parameters were empirically defined for clean detection of larval movement with minimal noise. PTZ was diluted in sterile distilled water and either PTZ or water alone was pipetted directly into each well and the assay proceeded with the addition of PTZ at 10mM. Activity data from pre- and post-PTZ were visualized in line graphs. Data were analyzed by normalizing the post-PTZ activity of each fish to its pre-PTZ values. Three-way LMM was used to tease apart gene, estradiol, and PTZ responses. The following parameters are generated by the Zebralab software to describe different levels of activity: frect (freeze count), fredur (freeze duration), midct (mid count), middur (mid duration), burct (burst count) burdur (burst duration), zerct (zero count), zerdur (zero duration), actinteg (action integration).

#### Bulk RNA extraction and sequencing

Total RNA was extracted from isolated larval heads with the eyes removed (n = 3 heads per genotype/condition, 1 head per sample), as described in^38^. To control for genetic background, RNA was extracted from sibling- or cousin-matched larvae from heterozygous incrosses of *scn1lab^Δ44/+^* fish exposed to 20 µM estradiol dissolved in DMSO or DMSO alone for 24 h from 5-6 dpf. Larvae were genotyped at 3 dpf using the Zebrafish Embryo Genotyper (ZEG) system (wFluidx, Salt Lake City, UT, USA) as previously described^38^. At 6dpf, larvae were euthanized on ice for 15 minutes, after which the heads were isolated, and the eyes were removed using fine forceps. RNA was immediately extracted using the RNeasy Micro Kit (Qiagen). RNA library preparation was performed using NEBNext Single Cell Low Input RNA Library Prep Kit (Cat# E6420L) and sequenced at 25 million reads/sample using NovaSeq X Plus (Illumina) (Yale Center for Genome Analysis).

#### Identification of differentially expressed genes from bulk RNA-seq data

To identify DEGs, low quality reads were trimmed and adaptor contamination was removed using Trim Galore (v0.5.0 https://www.bioinformatics.babraham.ac.uk/projects/trim_galore/). Trimmed reads were mapped to the *Danio rerio* reference genome (GRCz11) using HISAT2 (v2.1.0)^149^. Gene expression levels were quantified using StringTie (v1.3.3b)^150^with gene models (release 103) from Ensembl. Differential expression analysis was conducted using DESeq2 (v 1.22.1)^151^.

#### Bulk RNA-seq pathway analysis

To understand the biological functions of rescued DEGs, we performed functional pathway analysis using PANTHER ^152^. We first mapped zebrafish genes to human orthologs using EnsemblBioMart (version GRCz10/danRer10)^153^, as previously described^38^. To identify pathways reversed following exposure to estradiol, we first identified DEGs in mutant + no E2 versus wild-type, using a nominal p < 0.1 and DEGs in mutant + estradiol vs wild-type, using a nominal p <0.1. We then categorized DEGs from both comparisons into upregulated (fold change > 1) and downregulated (fold change < 1) groups and ran PANTHER to identify significant biological pathways (FDR <0.05) from Gene Ontology (GO) biological processes, cellular components, and molecular functions. We used InteractiVenn to explore pathway intersections of [mutant+ DMSO downregulated compared to mutant + estradiol upregulated] and [mutant + DMSO upregulated compared to mutant + estradiol downregulated] to determine estradiol reversed pathways.

##### Meta-analysis integration of multi-modal transcriptomic, cellular, and behavioral datasets

The single-cell RNA data, exists as present as n cells and m genes, whereas calcium imaging data is present as n cells and m phenotypes. On the other hand, zebrafish data present as n cells and m phenotypes (stimulated and awake at different conditions). The three modalities: scRNA, Calcium imaging, and zebrafish data exist in their own respective spaces. With the goal of uncovering the treatment effect of E2 across all three modalities, we initially assume each modality as its own unique Metric Measure Space, given as:

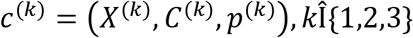

Where:

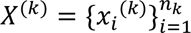 : cells in modality k

*C*^(*k*)^Î*R^n^k x nk* : intra-modality distance space (or pair-wise distances)

*p*^(*k*)^ = Δ*nk* : probability measure over cells

With the help of Metric Measure Space (MMS), the relative geometry (pair-wise relationship) of cells within each modality is determined by intra-modality distance, i.e., distance between each cell in their own modality, resulting in pair-wise distances in each respective modalities, with uniform weights to each cell, i.e. (*p*^(*k*)^) is a uniform distribution.

In the next step, these pair-wise distances (or the relational structure matrix) in different modalities are matched together using Gromov-Wasserstein (GW) distance framework, which essentially matches analogous relationships across modalities. For example, the GW distance between two MMS’s (*C*^(*a*)^, *p*^(*a*)^) and (*C*^(*b*)^, *p*^(*b*)^) is:

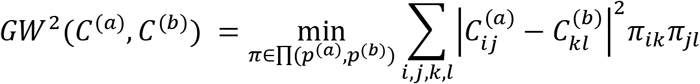

where:

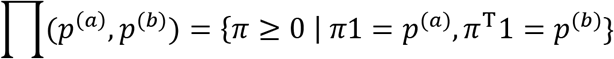

*π* is the transport matrix, giving probability of every pair of cells across modality, i.e. how likely *x_i_* ⟷ *y_k_* where *x_i_* is a point from modality 1, and *y_k_*is a point from modality 2.

Once the intra-modal relational structure and it’s cross-modal analogs are defined, we define barycenters of synthetic center of masses which represent cells across all modalities. Such barycenters are defined as having a structure:

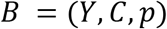

Where:

Y : synthetic cells (barycenters)

CÎ*R^N^ ^x^ ^N^*: barycenter distance matrix

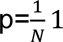 : relative probabilities

To clearly define these barycenters, a new shared latent space across modalities (*C*^∗^) is created, which needs to be solved by minimizing relative distances between barycenters against the distance between points in the modalities,

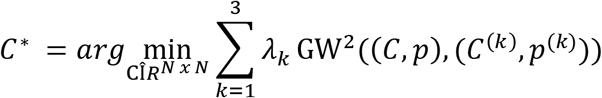

Which can also be written as:

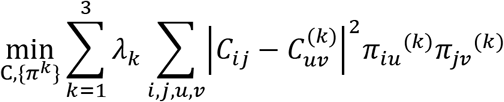

The coupling matrices are subjected to probability product:

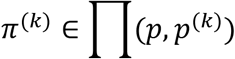

Based on these this, Optimal Transport (OT) framework is used to solve the barycenter, for which, the minimizer alternates between:

**GW alignment step:**

For fixed C, solving:

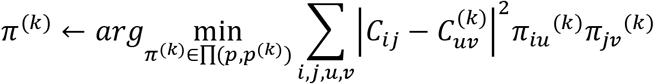

And **Barycenter update**:

For fixed *π*^(*k*)^,

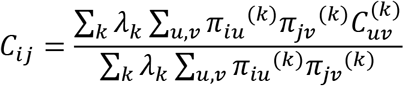

Using this solver, the optimization is solved via an alternating scheme, that iteratively updates GW couplings for fixed C, and then updates the barycenter distance matrix using a closed-form expression under squared loss. Once the shared latent space is learned, classical Multi-dimensional scaling (MDS) is applied to the distance matrix using eigen decomposition to generate low-dimensional Euclidean embeddings for further analysis.

##### Comparison of most and least estradiol-responsive genes

*De novo* protein-truncating variants (PTV) variants from 34,471 individuals with ASD across SPARK, ASC, SSC and MSSNG, together with 31,052 individuals with developmental disorders (DD) from DDD13K, GeneDx and RUMC76 were separately analyzed in the most (n = 7, *ASH1L, SCN2A, SHANK3, ASXL3, MED13L, CACNA1G, ANK2*) and least (n = 29, *ANK3, ARID1B, BCL11A, CHD2, CHD8, CREBBP, CUL3, DPYSL2, DYRK1A, FOXP2, GRIA3, GRIN2A, KDM5B, KDM6B, KMT2C, KMT5B, MBD5, NRXN1, PHF12, PHF21A, POGZ, PPP2R5D, PTEN, SETD5, SIN3A, SKI, SLC6A1, SMARCC2, WAC*) estradiol-responsive ASD/NDD genes. Clinical phenotypes across six standardized phenotypes (FSIQ, SCQ, DCDQ, RBSR, SRS, and VABS) were compared between male and female participants in SPARK with variants in the seven most-and 29 least estradiol-responsive ASD/NDD genes, using PTV-only and PTV+MisB sensitivity analysis.

Using differential expression results from the UCLA and BrainVar cortical transcriptomic datasets and their meta-analysis^77^, we tested association with baseline sex-biased cortical expression by comparing the absolute male–female log2 fold change between the two gene groups. Developmental-stage-specific co-expression was examined using BrainSpan^154^ cerebral cortex transcriptomic data. Gene–gene Spearman correlations were calculated separately across seven developmental stages. For each gene, within-group connectivity was defined as its mean Spearman correlation with the other genes in the same group. Gene-wise standardized expression across was compared across 16 cell types in the developing human cortex^78^. Within-group PPI connectivity was assessed using the STRING functional interaction network^60^. Because the two gene groups differed substantially in size, the degree of each gene was normalized by the maximum possible number of within-group interactions using Wilcoxon rank-sum test.

## STATEMENT OF ETHICS

Yale University Institutional Review Board waived ethical approval for this work. Ethical approval was not required because the hiPSC lines, lacking association with any identifying information and widely accessible from a public repository, are thus not considered to be human subject research. Post-mortem brain data are similarly lacking identifiable information and are not considered human subject research.

All procedures involving zebrafish were conducted in accordance with Institutional Animal Care and Use Committee (IACUC; Protocol #2024-20054) regulatory standards at Yale University.

## AUTHOR CONTRIBUTIONS

AP designed and executed the NDD gene ECCITEseq in NPCs, with mentorship from MFG and technical assistance from AS. NB designed and executed the NDD gene ECCITEseq in iGLUTs, with technical assistance from JL and IW. SL conducted all ECCITEseq bioinformatic analyses, with assistance from CS, AS, KRT, XL, and HG. Arrayed KO lines and neuronal cultures were generated, validated, phenotyped, and analyzed by NB. LY conducted and analyzed arrayed calcium imaging experiments, performed cross-modality meta-analysis, and iGLUT MEA. Cell culture support provided by PJMD. Zebrafish mutations were generated, phenotyped, and analyzed by AP, ED, YD, SK, and MC with technical assistance from GD, bioinformatic assistance from GX and XH, and bioinformatic supervision from ZW and XP. AP performed and analyzed zebrafish RNAseq, PTZ seizures, and whole-brain activity mapping experiments, with assistance from PJ and bioinformatics analysis support from DZ. S.G. and T.W. conducted post-hoc comparisons of estradiol responsive and non-responsive gene sets. Funding and mentorship provided by LH, EJH, and KJB. Manuscript was written by AP and KJB with extensive feedback from LY, SL, NB, LH, and EJH, as well as contributions from all authors.

Special thanks to Michael Talkowski for CRISPR-engineered *SCN2A* hiPSC lines and to Summer Thyme and the Thyme lab as well as Julia Dallman and the Dallman lab for sharing zebrafish mutant lines.

## DATA AVAILABILITY

All source donor hiPSCs have been deposited at the Rutgers University Cell and DNA Repository (study 160; http://www.nimhstemcells.org/). sc-RNA sequencing data reported in this paper will be uploaded to Gene expression omnibus (GEO) prior to publication. Previously published SCZ-CRISPRa screen datasets that were used for external validation of random forest models are available on the GEO (GSE200774) and on Synapse (syn27819129).

Zebrafish bulk RNA sequencing data reported in this paper will be uploaded to Gene expression omnibus (GEO) prior to publication.

## CODE AVAILABILITY

The full analysis pipeline (including code and processed data objects) used for analysis of single-cell CRISPR-KO data, evaluation and characterization of gene-level and network level convergence, and predictive modeling using random forest will be publicly available through Synapse prior to publication.

Custom MATLAB software developed by the Hoffman Lab to analyze visual-startle response parameters is available on github at https://github.com/ehoffmanlab/Weinschutz-Mendes-et-al-2023-behavior; https://doi.org/10.5281/zenodo.7644898. Custom MATLAB software developed by Jason Rihel to analyze sleep-wake assays is available on github at https://github.com/JRihel/Sleep-Analysis/tree/Sleep-Analysis-Code; https://doi.org/10.5281/zenodo.7644073.

