## Supplementary material for "Convergent and divergent phenotypic impacts of estradiol on neurodevelopmental disorder risk genes": SI Figures

INVENTORY OF SUPPLEMENTAL MATERIAL

*SI Figure 1: Estradiol rescues key molecular pathways in NPCs.*

*SI Figure 2: Expression levels of ESR1, ESR2, and GPER in neural cells.*

*SI Figure 3: Estradiol dose response in iGLUTs and larval zebrafish.*

*SI Figure 4: Validation of gRNAs in estradiol ECCITE-seq data.*

*SI Figure 5: Degree of estradiol rescue in ASH1L and SCN2A iGLUTs.*

*SI Figure 6: Convergent and rescued DEGs are not more likely to be ER targets.*

*SI Figure 7: Feature-based clustering of immature network calcium activity.*

*SI Figure 8: Clustering of mature network calcium oscillation patterns across NDD mutants.*

*SI Figure 9: Sensorimotor and arousal behavioral screen identifies genes susceptible to estradiol rescue.*

*SI Figure 10: Estradiol differentially rescues behavioral phenotypes across 4 NDD mutants.*

*SI Figure 11: Estradiol rescues dysregulated molecular pathways.*

*SI Figure 12: Effect of estradiol on wild-type tERK intensity.*

*SI Figure 13: Effect of estradiol on whole brain activity in scn1lab$^{-/-}$ mutants.*

*SI Figure 14: Effect of estrogen receptor agonists on seizure behavior in scn1lab$^{-/-}$ mutants.*

*SI Figure 15: Sex-biased genetic and clinical effects of estradiol-responsive ASD/NDD genes.*

*SI Figure 16: Degree of rescue thresholds criteria.*

*SI Table 1: ASD/NDD risk genes functional classification*

*SI Table 2: NPC 29 gene library*

*SI Table 3: iGLUT 36 gene library*

*SI Table 4: Final number of cells in NPC and iGLUT*

*SI Table 5: iGLUT E2  Rescue pathways*

*SI Table 6: NPC E2 Rescue pathways*

*SI Table 7: Zebrafish Behavioral glossary*

*SI Table 8: Zebrafish F0 CRISPR gRNAs*

*SI Table 9: Zebrafish stable mutant line CRISPR gRNAs*

*SI Table 10: Zebrafish baseline behavior p-value scores*

*SI Table 11: Zebrafish baseline behavior beta values*

*SI Table 12: Zebrafish E2 behavior scores*

*SI Table 13: Zebrafish ER Target Mechanism Enrichment*

*SI Table 14: Zebrafish Correlating and Anti-Correlating drug list*

*SI Table 15: scn1lab HOM DEGs*

*SI Table 16: scn1lab HOM rescued pathways*

*SI Table 17: scn1lab WT tERK intensity values*

*SI Table 18: Brain Activity Mapping Regions by Degree of Rescue*

*SI Table 19: p-values for seizure LMM*

*SI Table 20: p-values for E2 MEA*

*SI Table 21: Overview of E2 gene-x-modality rescue*

*SI Table 22: Ranking of KO genes by E2 rescue effect*

*SI Table 23: Cohorts for de novo variant analysis*

*SI Table 24: Sex-stratified PTVs in ASD*

*SI Table 25: Sex-by-gene interactions*

*SI Table 26: Gene co-expression connectivity across development*

*SI Table 27: Developmental expression profiles*

*SI Table 28: Cell type expression by E2 responsive group*

*SI Table 29: Cortical cell type expression profiles*

*SI Table 30: Sex-biased cortical expression*

*SI Table 31: iGLUT convergent pathways across KO effect*

*SI Table 32: Percent convergent genes across KOs*

*SI Table 33: Recipes for NIM and NMM*

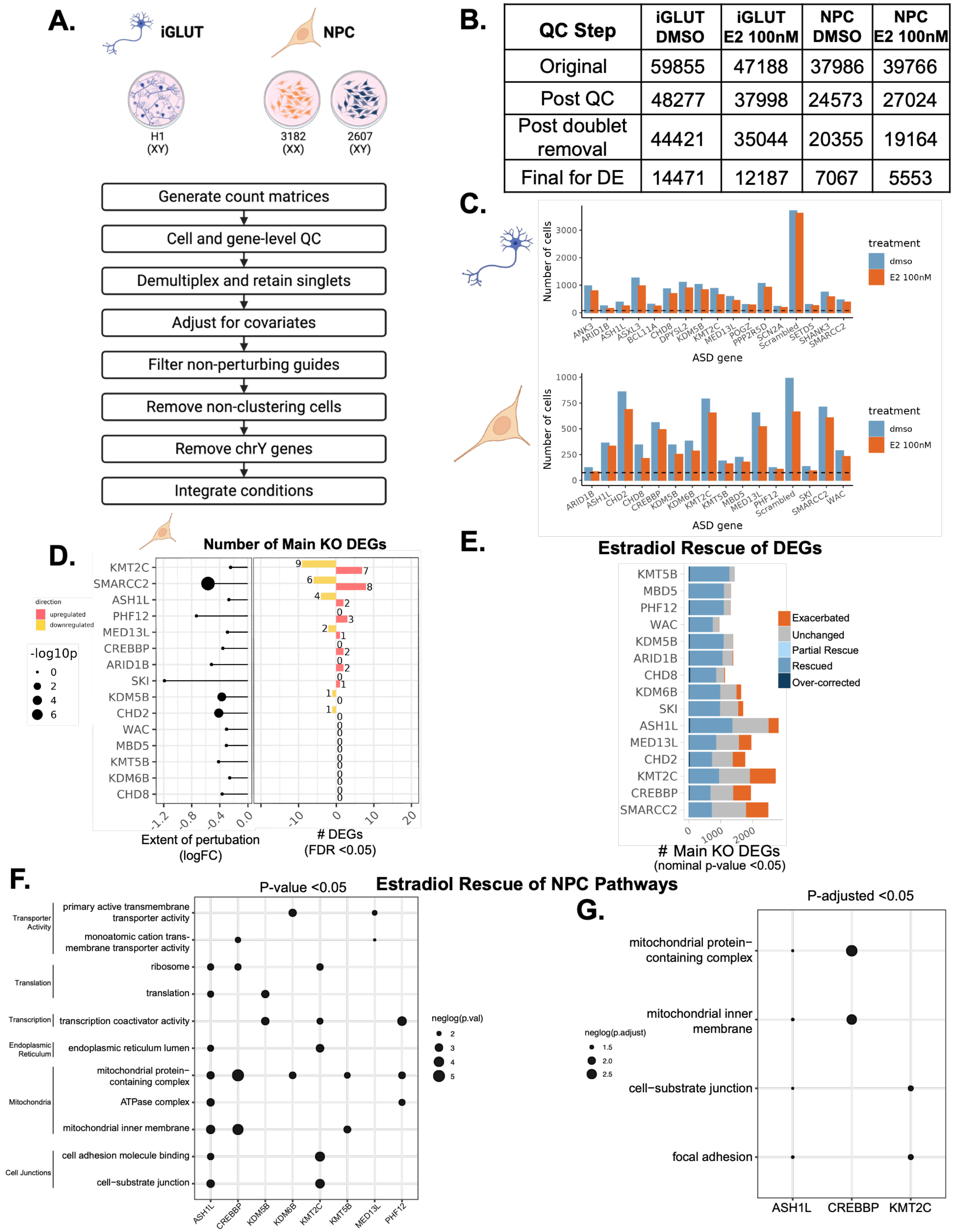


| QC Step | iGLUT DMSO | iGLUT E2 100nM | NPC DMSO | NPC E2 100nM |
|---|---|---|---|---|
| Original | 59855 | 47188 | 37986 | 39766 |
| Post QC | 48277 | 37998 | 24573 | 27024 |
| Post doublet removal | 44421 | 35044 | 20355 | 19164 |
| Final for DE | 14471 | 12187 | 7067 | 5553 |

***SI Figure 1: Estradiol rescues key molecular pathways in NPCs.*** **A**. Experimental design and analysis pipeline post sequencing. **B.** Number of cells at each step of quality control. **C.** Final number of cells per target identity by cell type and treatment condition (top, iGLUTs; bottom, NPCs). **D.** Extent of perturbation per target NDD gene (left, dot plot). Circle size indicates negative $\log_{10}$(p-value). Stem indicates the log fold change of the knockout +DMSO vs non-targeting control + DMSO. KOs were called based on enrichment of gRNA for target NDD genes and downregulation of target KO gene expression. Number of differentially expressed genes in the main KO effect are shown in the bar plot (FDR <0.05). **E.** Effect of estradiol rescue on DEGs across 16 target NDD genes knockouts in NPCs. Target genes are ranked from highest to lowest percentage of DEGs that are classified as rescued (nominal p-value <0.05). **F-G.** Convergent estradiol rescue of biological pathways across NDD targets. Dot size indicates the –log10 of the gene set enrichment significance of E2 effect within KO at p-value <0.05 **(F)** and at p-adjusted value <0.05 **(G).**

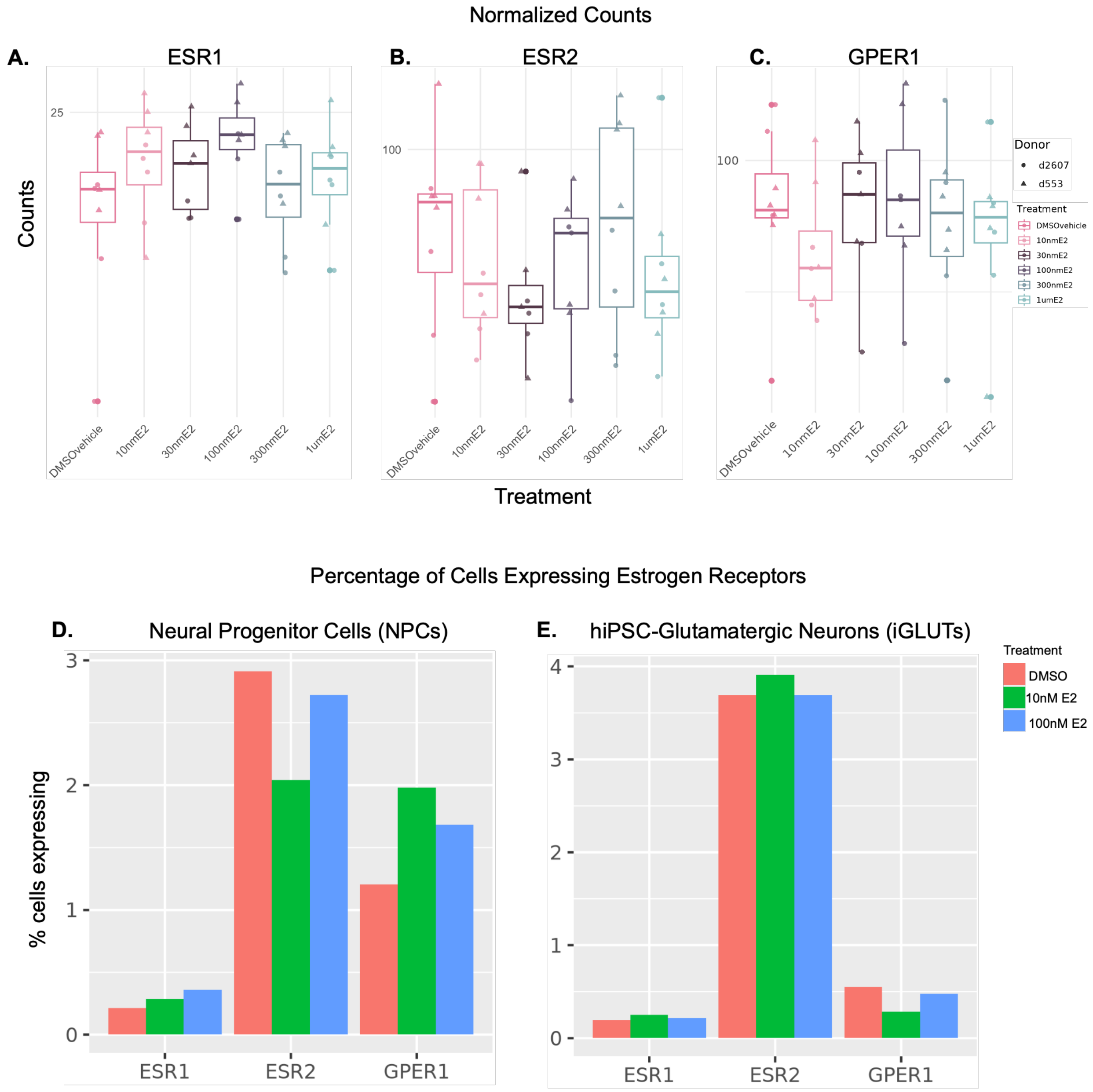


***SI Figure 2: Expression levels of ESR1, ESR2, and GPER in neural cells.*** Expression of the nuclear (*ESR1, ESR2)* and membrane (*GPER)* estrogen receptor in control neural cell types. **A-C.** Dose response bulk RNAseq normalized counts of estrogen receptors (ERs) *ESR1, ESR2,* and *GPER1* in iGLUTs. Shapes of data points indicate two individual control male donors (d2607 and d553). Doses of estradiol (E2): 10nM, 30nM, 100nM, 300nM, and 1$\mu$M or DMSO vehicle. **D-E.** Percentage of cells expressing ERs at 10nM E2, 100nM E2, and DMSO vehicle in neural progenitor cells (NPCs) **(D)** and iGLUTs **(E).**

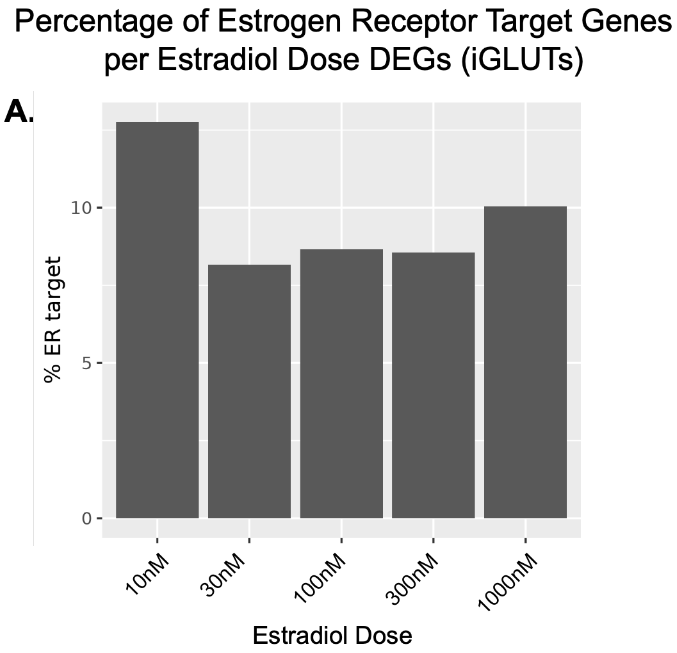


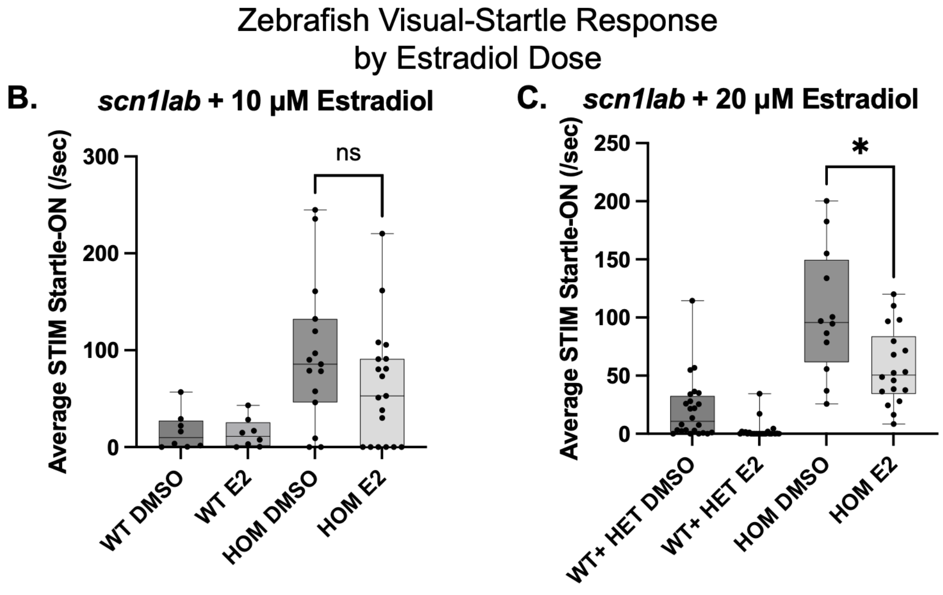


***SI Figure 3: Estradiol dose response in iGLUTs and larval zebrafish.*** **A.** Percentage of estrogen receptor target genes per estradiol dose DEGs from bulk RNAseq data in iGLUTs from two individual control male donors (d2607 and d553). Doses of estradiol (E2): 10nM, 30nM, 100nM, 300nM, and 1000nM. Higher doses have moderate percentage of ER target DEGs resulting from 24hr estradiol treatment. **B-C.** Measurement of the average stimulus (STIM) startle response to lights-ON in *scn1lab* homozygous, heterozygous, and background matched wild-type with DMSO vehicle or estradiol at 10$\mu$M (**B**) and 20 $\mu$M (**C**). STIM response was significantly decreased with 20 μM treatment. Sample sizes: for 10$\mu$M, WT DMSO n=8, WT E2 n=8, HOM DMSO n=15, HOM E2 n=19. For 20 $\mu$M, WT/HET DMSO n=24, WT/HET E2 n=24, HOM DMSO n=12, HOM E2 n=18.

**A.**

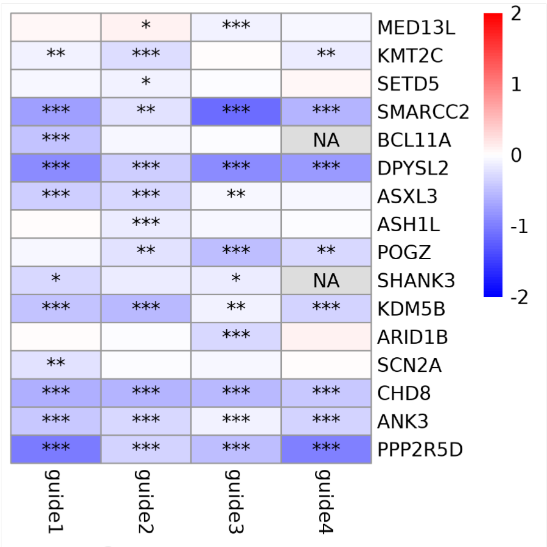


**B.**

| Target Gene | Min Cor | Max Cor |
|---|---|---|
| ANK3 | 0.70 | 0.83 |
| ASXL3 | 0.70 | 0.79 |
| CHD8 | 0.69 | 0.73 |
| DPYSL2 | 0.75 | 0.85 |
| KDM5B | 0.72 | 0.83 |
| KMT2C | 0.66 | 0.81 |
| POGZ | 0.79 | 0.79 |
| PPP2R5D | 0.74 | 0.87 |
| SHANK3 | 0.73 | 0.73 |
| SMARCC2 | 0.80 | 0.83 |

**C.**

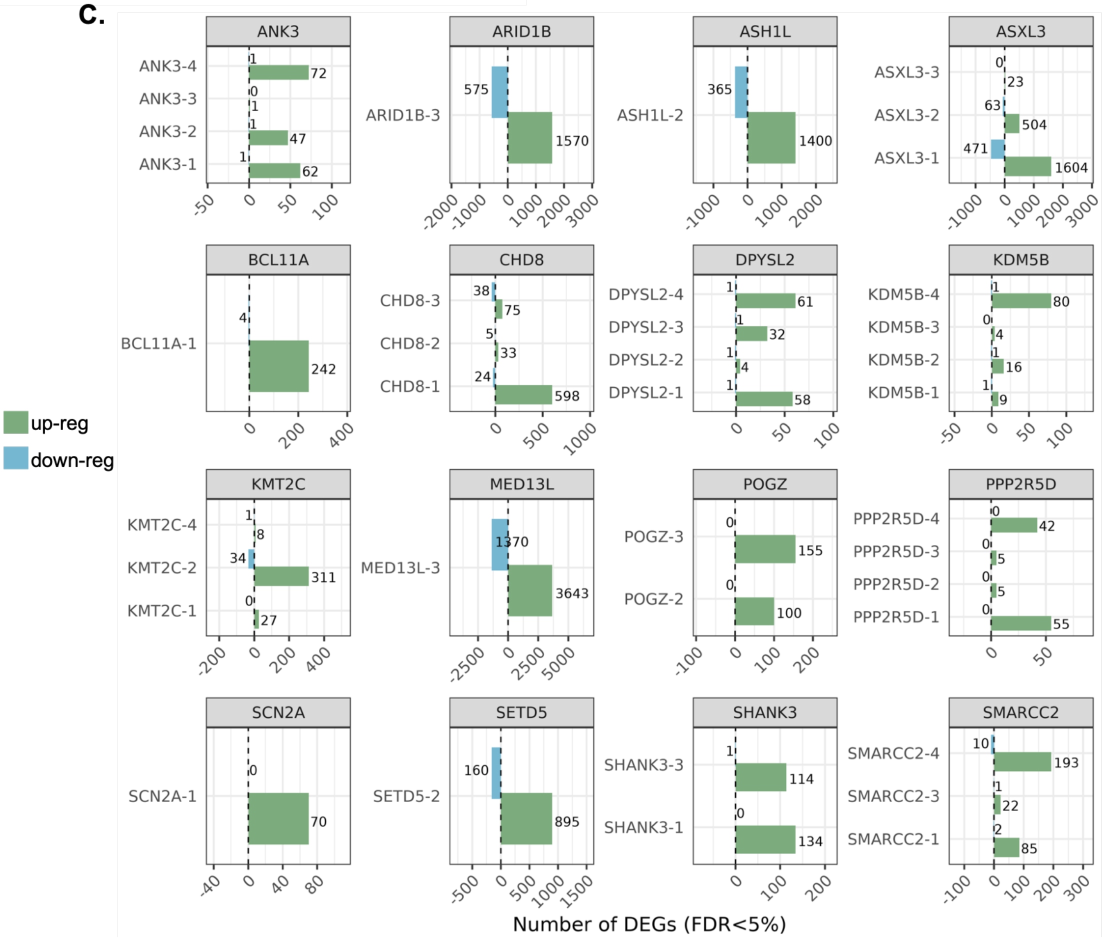


***SI Figure 4: Validation of gRNAs in estradiol ECCITE-seq data.*** **A.** Average logFC of each gRNA vs Scrambled cells, per ASD/NDD target gene. Asterisks indicate ***p<0.01, **p<0.05, *p<0.1. **B.** Table of minimum and maximum Pearson correlations between gRNAs per target gene. **C.** Number of DEGs per gRNA for each target gene. FDR <5%.

### ASH1L

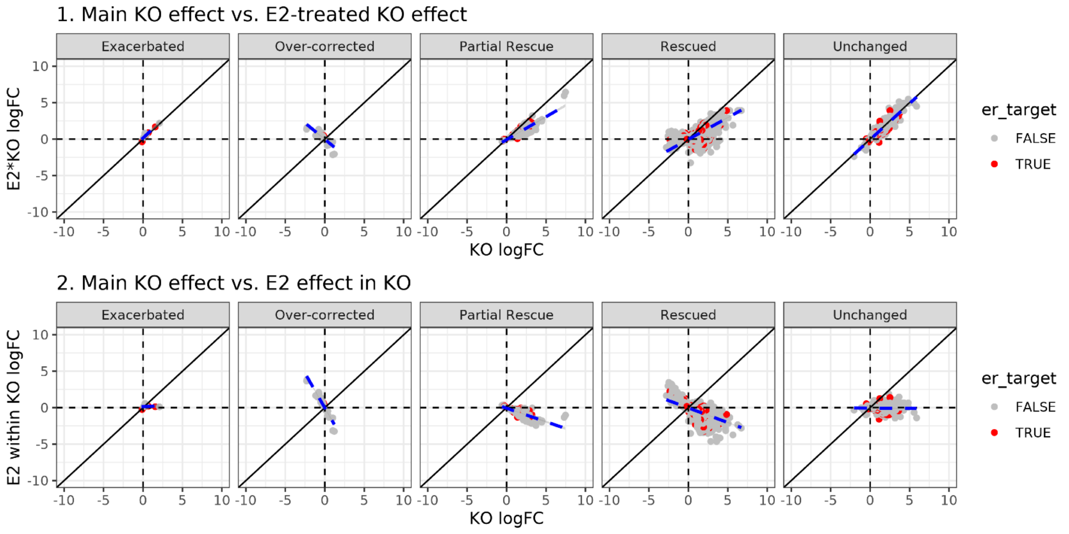


### SCN2A

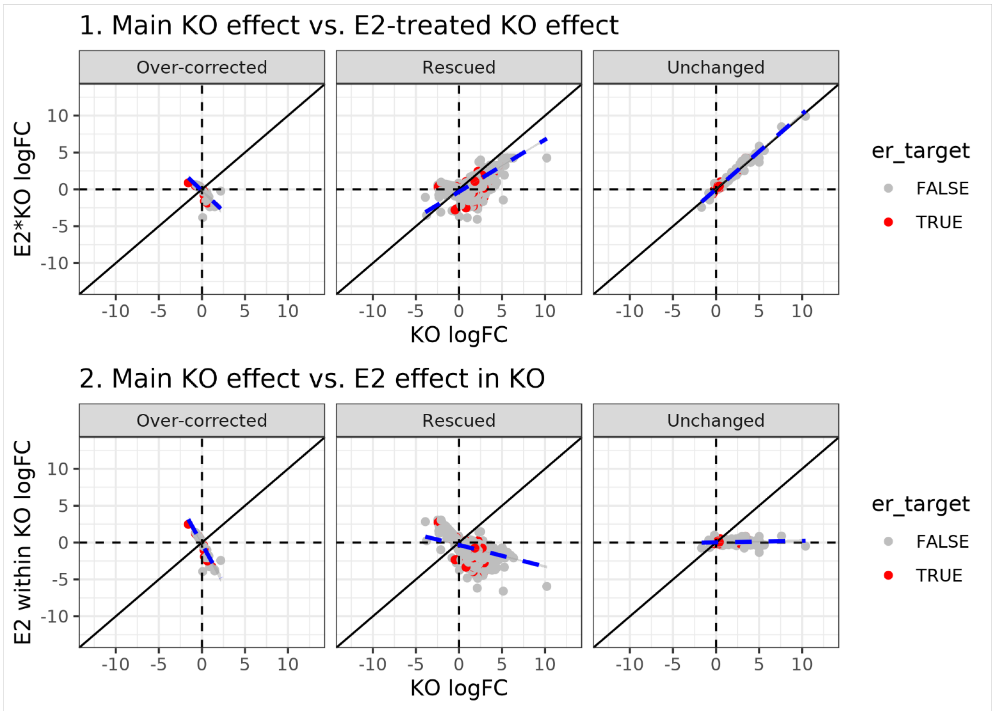


***SI Figure 5: Degree of estradiol rescue in ASH1L and SCN2A iGLUTs.*** For two top ASD/NDD target genes, *ASH1L* and *SCN2A*, panel 1 (top) compares the main KO effect vs the estradiol treated KO effect. Here, for the *partial rescue* and *rescue* categories, the blue line is angled to the right compared to the reference line (black line), indicating the effect sizes of the main KO genes were ameliorated with E2 treatment. Panel 2 (bottom) compares the main KO effect vs estradiol effect in KO. Here, there is a negative correlation of the KO and estradiol effects.

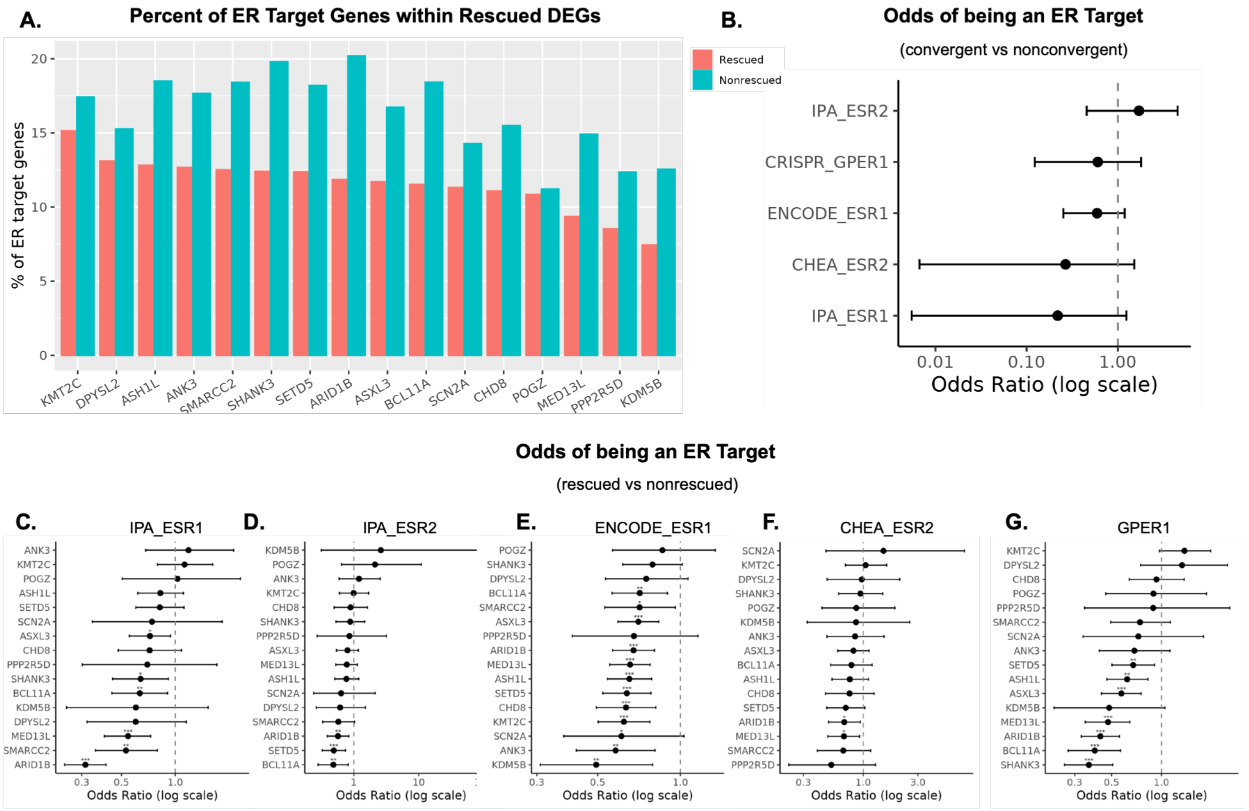


***SI Figure 6: Convergent and rescued DEGs are not more likely to be ER targets.*** **A.** Percentage of ER target genes (ESR1/ESR2/GPER1) in rescued or non-rescued DEGs. Target NDD KOs are ranked by percentage of ER target genes rescued. **B.** Odds of convergent genes being an ER target compared to nonconvergent genes, tested across individual ER target gene sets. **C-G.** Odds of rescued genes being an ER target compared to non-rescued genes for *ESR1* **(C,D),** *ESR2* (**E,F),** and *GPER1* **(G).** Fisher's exact test, ***p<0.001, **p<0.01, *p<0.05. Gene sets: IPA, Ingenuity Pathway Analysis; ENCODE, ENCODE Transcription Factor Targets; CHEA, ChEA Transcription Factor Targets; GPER1, LINCS L1000 CMAP CRISPR Knockout Consensus Signatures. Gene sets can be accessed from Harmonizome 3.0 (https://maayanlab.cloud/Harmonizome/).

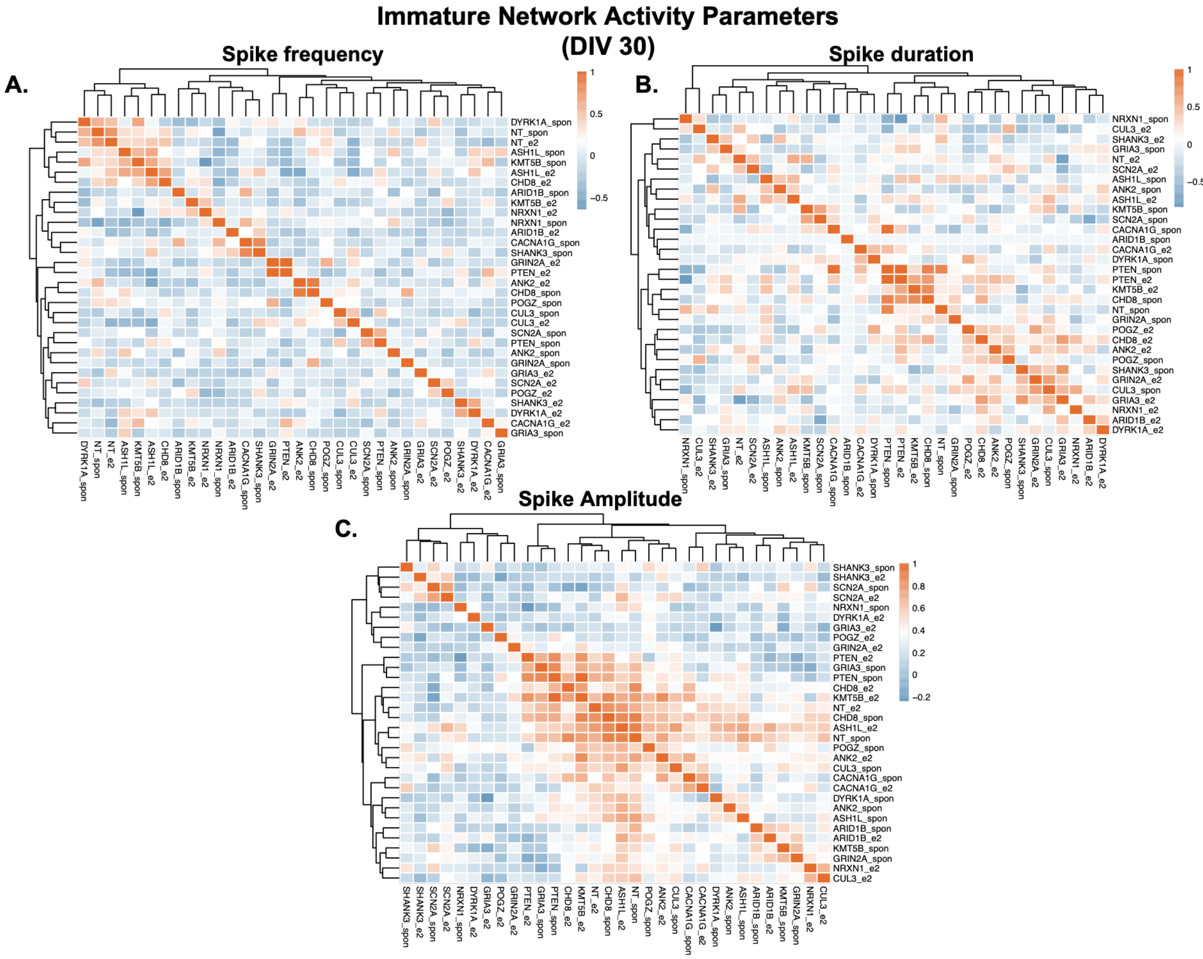


***SI Figure 7: Feature-based clustering of immature network calcium activity.*** Heatmaps depict pairwise similarity of spike frequency **(A),** duration **(B),** and amplitude **(C),** across all KOs and control conditions under spontaneous and estradiol-treated states at 30 DIV. Rows and columns correspond to perturbation–condition pairs, with hierarchical clustering performed independently for each feature. Color scales indicate normalized similarity values, and dendrograms reflect unsupervised clustering performed independently for each calcium activity feature.

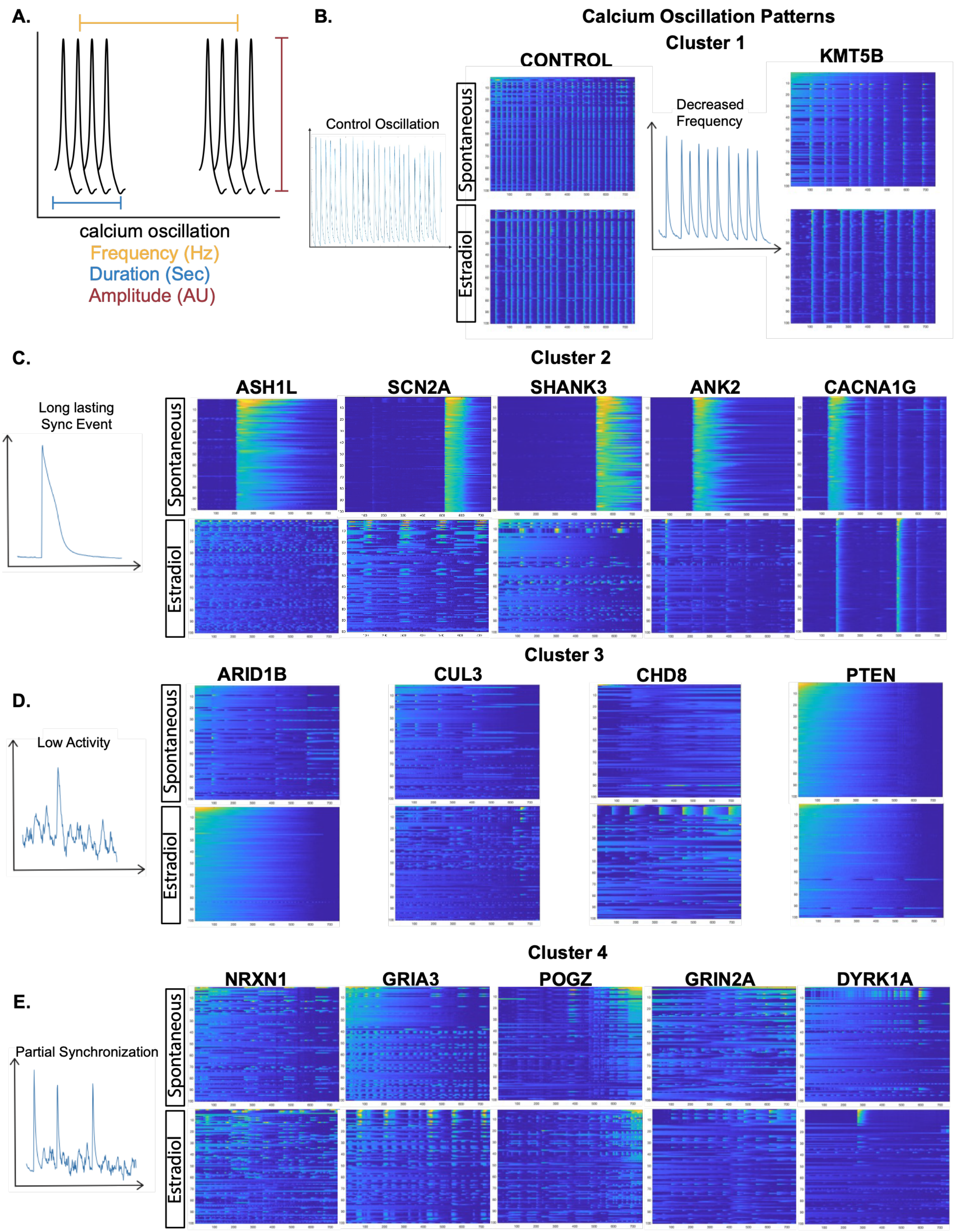


***SI Figure 8: Clustering of mature network calcium oscillation patterns across NDD mutants.*** **A.** Schematic illustrating calcium oscillation features quantified in this study, including spike frequency, duration, and amplitude. **B-E.** Raster plots show calcium activity traces for control and KO conditions under spontaneous and estradiol-treated states at 60 DIV, grouped based on oscillation patterns. Cluster 1 is characterized by reduced oscillation frequency relative to control (*KMT5B*); Cluster 2 by long-lasting synchronized events (*ASH1L, SCN2A, SHANK3, ANK2, CACNA1G*), with altered patterns observed following E2 treatment; Cluster 3 by low overall activity (*ARID1B, CUL3, CHD8, PTEN*); and Cluster 4 by partially synchronized activity patterns (*NRXN1, GRIA3, POGZ, GRIN2A, DYRK1A*).

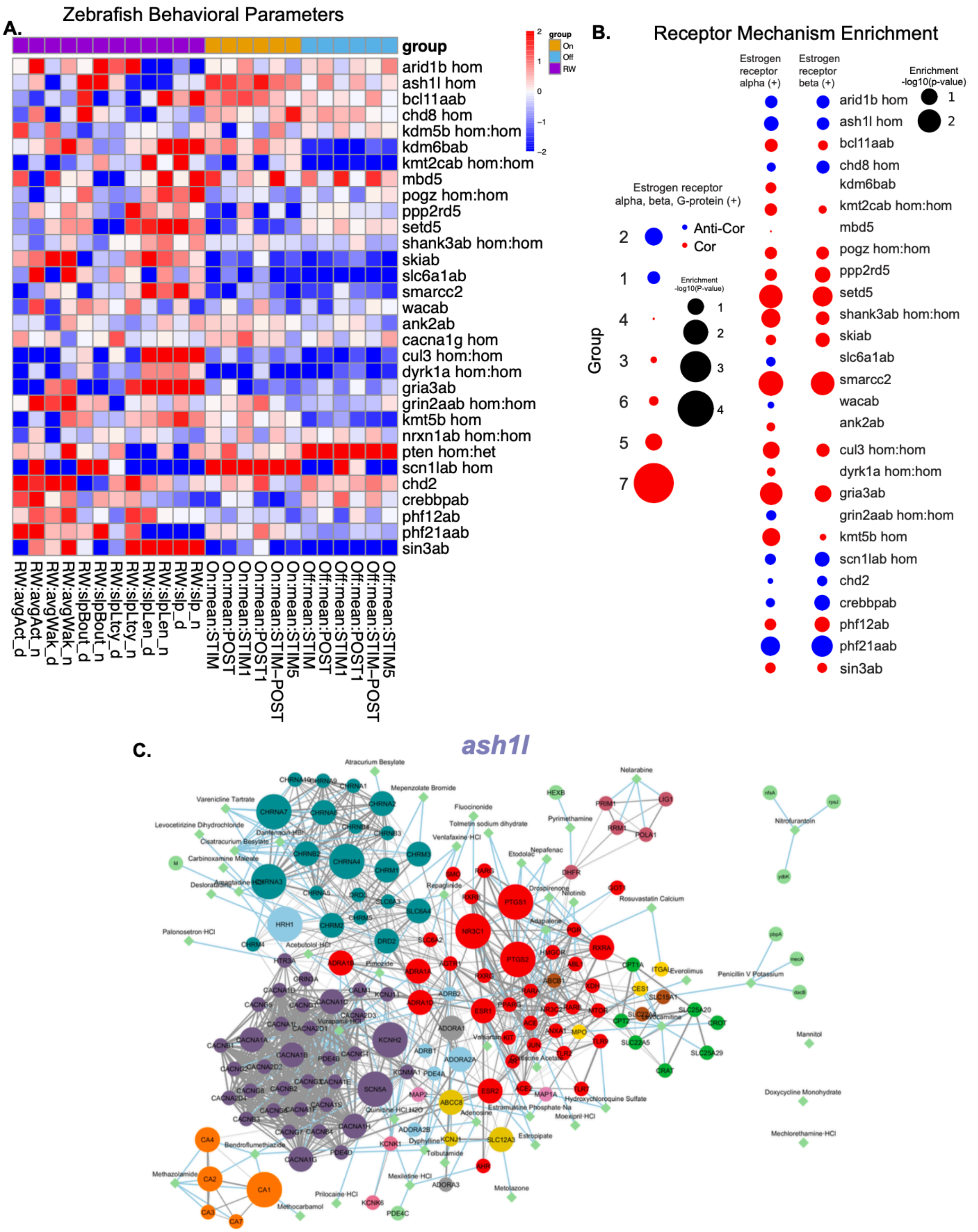


***SI Figure 9: Sensorimotor and arousal behavioral screen identifies genes susceptible to estradiol rescue.*** **A.** Hierarchical phenotypic clustering of behavioral fingerprints across 31 NDD genes in mutant zebrafish. Startle-OFF (light blue), startle-ON (orange), and sleep-wake (purple) parameters are shown. Each box represents the signed $-\log_{10}$-transformed p-values from linear mixed models (LMM) comparing behaviors in stable mutant and background-matched wild-type fish and F0 mutant and scrambled control (red, increased in mutant; blue, decreased in mutant). **B.** Enrichment of combined molecular targets *estrogen receptor (ER) alpha, beta, and G-protein* by behavioral subgroup (left) and separate ER-alpha and ER-beta by gene (right) among anti-correlating (blue, correlation <-0.5, p<0.05, t-statistic) and correlating drugs (red, correlation >0.5, p<0.05, t-statistic). Mutants for which there is no ER enrichment are not shown. Circle size indicates the $-\log_{10}$(p-value), calculated using Fisher's exact test. **C.** Extended protein-protein interaction networks among the targets (circles) of anti-correlating drugs (diamonds) (correlation <-0.5, p<0.05, t-statistic) for *ash1l*.

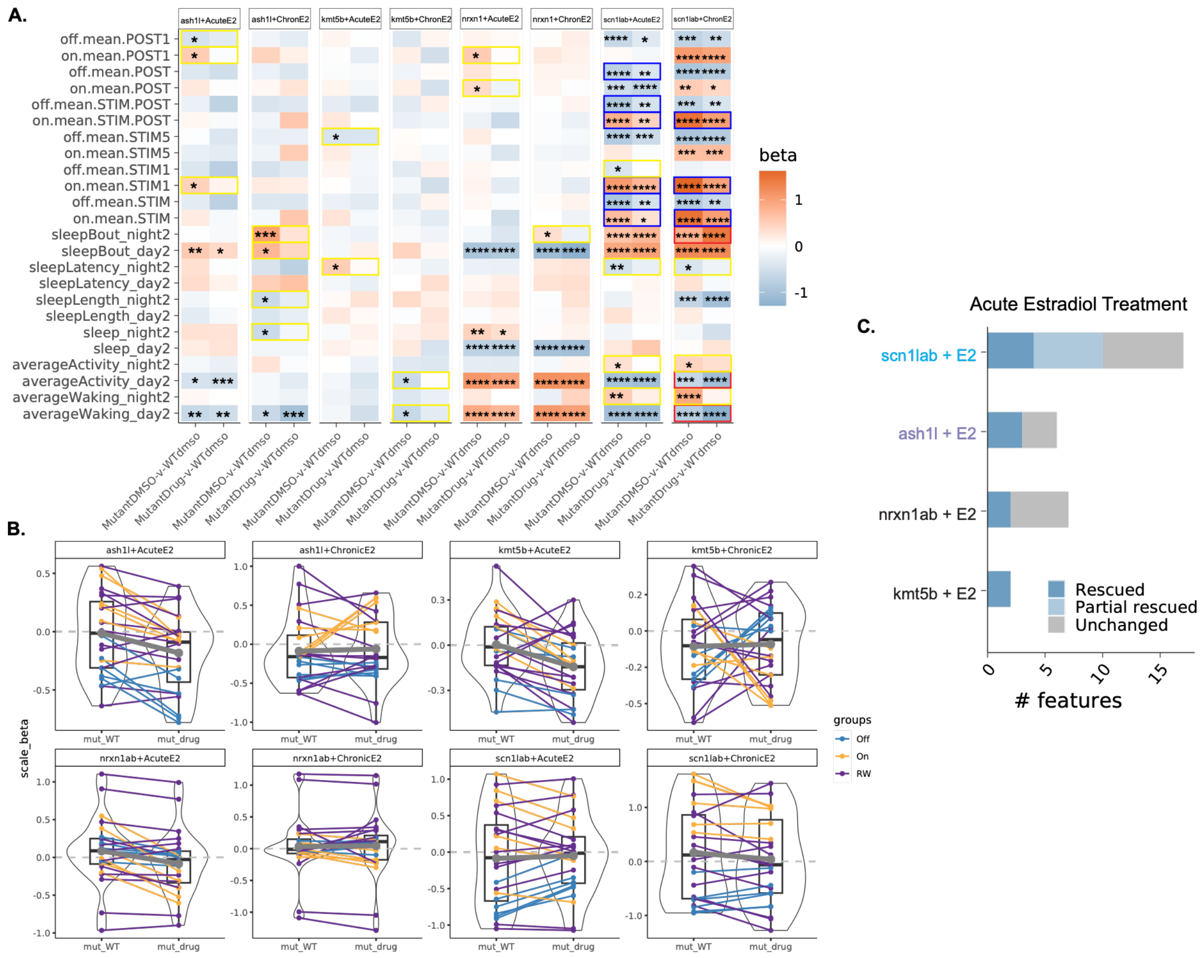


***SI Figure 10: Estradiol differentially rescues behavioral phenotypes across 4 NDD mutants.*** **A.** Heatmap of beta values for homozygous mutant + DMSO vs wild-type + DMSO (mutant effect alone) or mutant + E2 (acute or chronic) vs wild-type + DMSO (gene-x-E2 interaction effect) for each tested behavior. Color indicates beta values. Asterisks indicate significant differences in respective comparison group (p * <0.05, ** <0.01, ***<0.001, ****<0.0001, LMM). Rescued parameters in yellow, partially rescued parameters in dark blue, exacerbated parameters in red, unchanged parameters are indicated with asterisks without boxes. **B.** Comparison of the magnitude of effect (beta) on startle-ON (yellow), startle-OFF (blue), and sleep-wake (purple) behavioral parameters between [mutant+DMSO vs wild-type+DMSO] compared to [mutant+E2 vs wild-type+DMSO] groups (acute and chronic E2) for each empirical test. **C.** Ranking of estradiol rescue by mutant based on number of rescued and partially rescued features for acute estradiol treatment. Features were categorized as rescued, partially rescued, or unchanged.

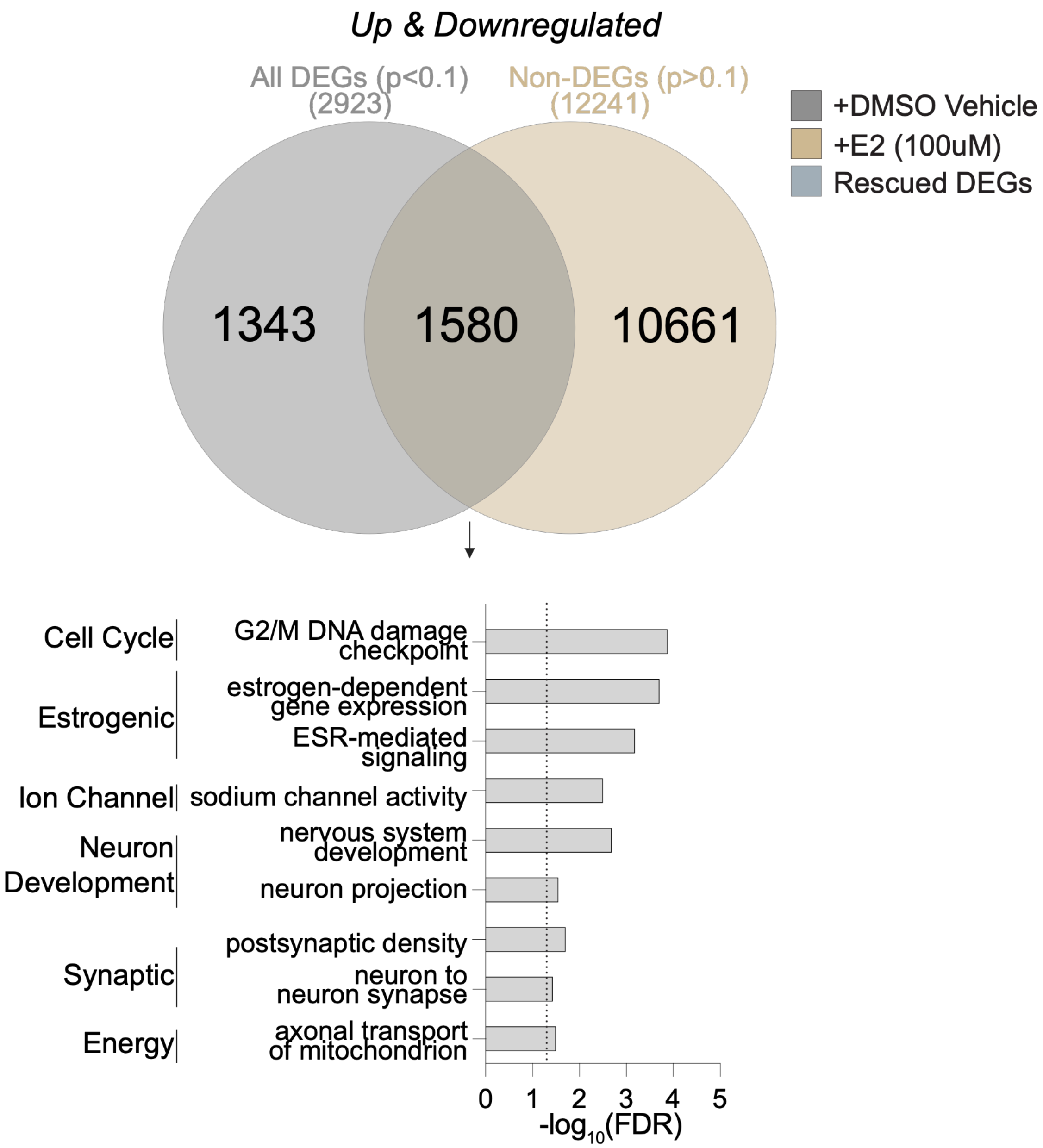


***SI Figure 11: Estradiol rescues dysregulated molecular pathways***. Venn diagram reflects the union of upregulated and downregulated conditions for all significant DEGs in the *scn1lab* homozygous + DMSO condition (left), non-DEGs following estradiol treatment (right) and rescued DEGs (middle). From the rescued DEGs, bar plot reflects Gene Ontology (GO)[1] pathways that were rescued in *scn1lab* homozygous mutants by estradiol treatment. For the full list of rescued pathways, see SI Table 16.

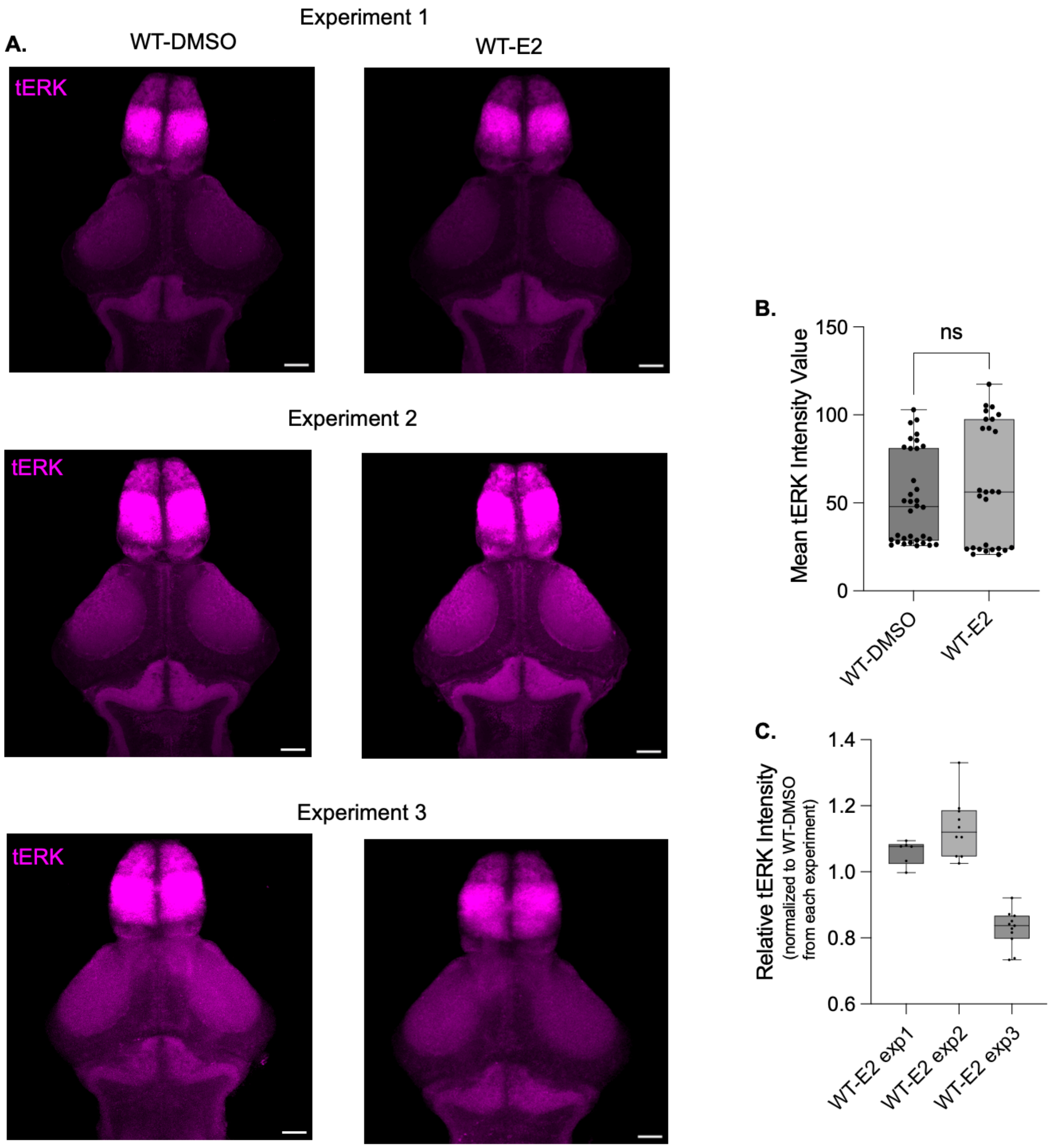


***SI Figure 12: Effect of estradiol on wild-type tERK intensity.*** **A.** Representative whole brain images of tERK stain in wild-type + DMSO condition (left) and wild-type + estradiol (E2, 20 $\mu$M) from three independent experiments. **B.** Average pixel intensity value for wild-type + DMSO and wild-type + E2. Images were analyzed in FIJI (ImageJ) by generating a maximum z-stack projection from the C1 channel of the .lif file, drawing ROI around the max projection brain, and using the *Analyze* > *Measure* function to acquire mean intensity values. Full quantitative values are included in SI Table 17. Unpaired t-test with Welch's correction. **C.** Relative tERK intensity by experiment. Wild-type + E2 sample mean intensity values were normalized to each experiment's WT+DMSO average intensity value.

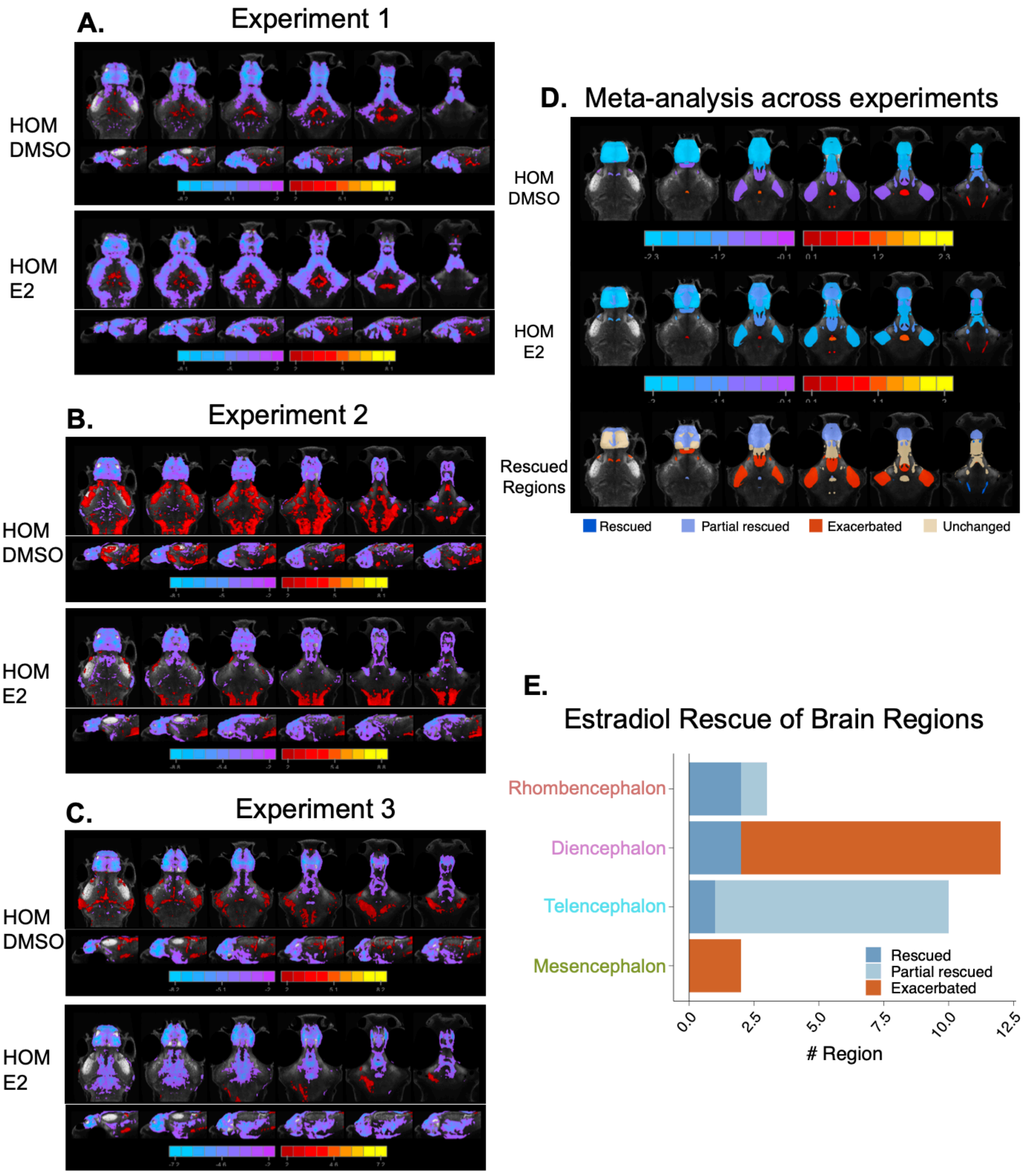


***SI Figure 13: Effect of estradiol on whole brain activity in scn1lab$^{-/-}$ mutants.*** **A-C.** Brain activity maps of voxel-wise Z score normalized pERK/tERK values for three independent experiments comparing *scn1lab* homozygous mutant + DMSO vs background matched wild-type + DMSO (top row) and *scn1lab* homozygous mutant + estradiol (20$\mu$M) vs background matched wild-type + DMSO (bottom row). Within the row, mosaic images are of the axial view (top, dorsal to ventral) and sagittal view (bottom, lateral to medial). Scale bar represents Z score (red/yellow, increased in mutant;

cyan/purple, decreased in mutant). **D.** Mosaic axial visualization of the meta-analysis across the 3 experiments for *scn1lab* homozygous mutant + DMSO vs background matched wild-type + DMSO (top) and *scn1lab* homozygous mutant + estradiol (20$\mu$M) vs background matched wild-type + DMSO (middle). Scale bar represents scaled beta values. Bottom row shows mosaic visualization of regions classified as exacerbated (orange), unchanged (beige), partial rescued (lavender), and rescued (dark blue). **E.** Bar plot of degree of rescue for number of regions in the 149 atlas by 4-region atlas.

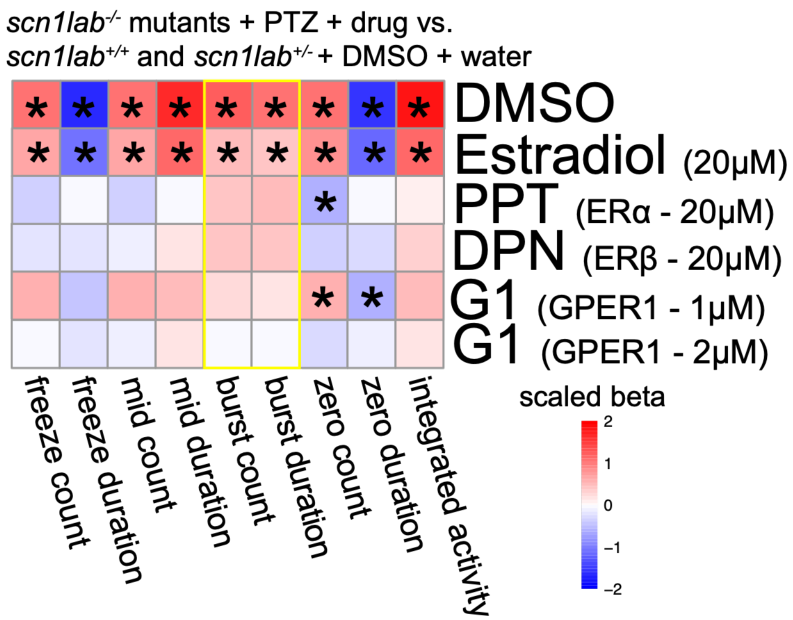


***SI Figure 14: Effect of estrogen receptor agonists on seizure behavior in scn1lab$^{-/-}$ mutants***. *scn1lab$^{-/-}$* mutants were exposed to DMSO vehicle, estradiol (20$\mu$M), ER$\alpha$ agonist PPT (20$\mu$M), ER$\beta$ agonist DPN (20$\mu$M), or GPER1 agonist G1 (1$\mu$M or 2$\mu$M) then treated with 10mM PTZ for the seizure assay. Heatmap shows scaled beta values from seizure parameters (x-axis) vs agonist treatment (y-axis). Asterisks indicate significant values by LMM. Each row represents *scn1lab$^{-/-}$* mutants treated with PTZ and the respective agonist or vehicle compared to background matched wild-type and heterozygous controls exposed to DMSO and treated with water. Data are shown from combining 4 independent experiments (estradiol treatment) and two independent experiments (PPT, DPN, G1). Yellow box highlights the burst count and burst duration parameters.

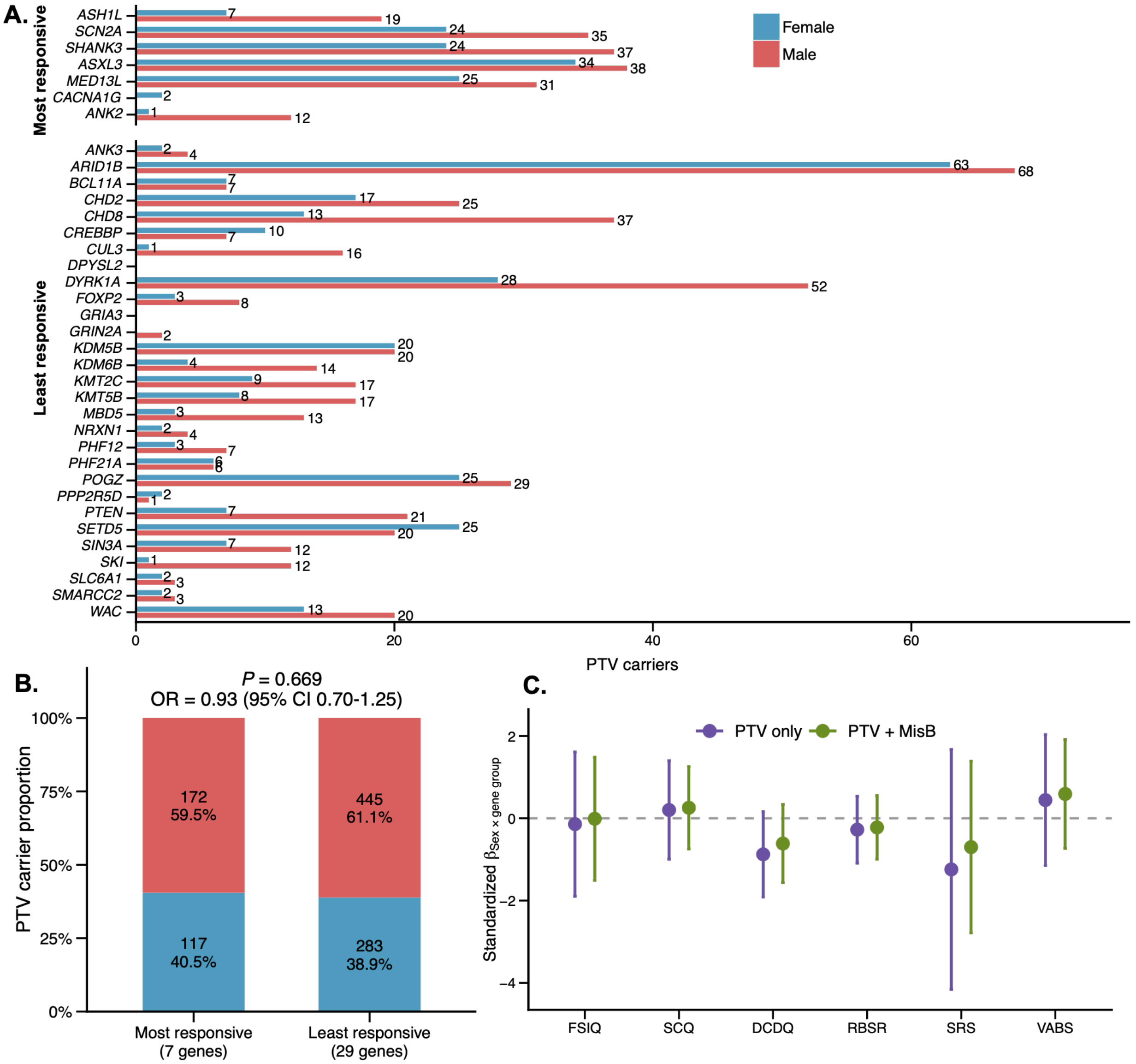


***SI Figure 15: Sex-biased genetic and clinical effects of estradiol-responsive ASD/NDD genes.*** **A,** Sex distribution of NDD probands carrying *de novo* protein-truncating variants (PTV) across the seven most-responsive and 29 least-responsive genes for estradiol. **B,** Comparison of the overall sex distribution of PTV carriers between the two gene groups (Fisher's exact test, OR = 0.93, 95% CI = 0.70–1.25, *P* = 0.669). **C,** Sex-by-gene-group interaction effects on six standardized ASD-related phenotypes in SPARK carriers: full-scale intelligence quotient (FSIQ), Social Communication Questionnaire (SCQ), Developmental Coordination Disorder Questionnaire (DCDQ), Repetitive Behavior Scale–Revised (RBSR), Social Responsiveness Scale (SRS) and Vineland Adaptive Behavior Scales (VABS). Points indicate standardized interaction coefficients (β) and error bars indicate 95% confidence intervals. Models were adjusted for phenotype-specific assessment age and the first 10 genetic principal components. PTV-only and PTV+MisB carrier definitions are shown separately in different color.

### Degree of Rescue Thresholds

$Y \sim \beta_0 + \beta_1 mutant + \beta_2 drug$

Mutant (or KO) + DMSO vs WT (or NT) + DMSO: $\beta_1$ ← “Mutant-WT” or “KO-WT”

Mutant (or KO) + Drug (E2) vs WT (or NT) + DMSO: $\beta_1 + \beta_2$ ← “Mutant-E2” or “KO-E2”

Mutant (or KO) + Drug (E2) vs Mutant (or KO) + DMSO: $\beta_2$ ← “E2-Effect”

Criteria:

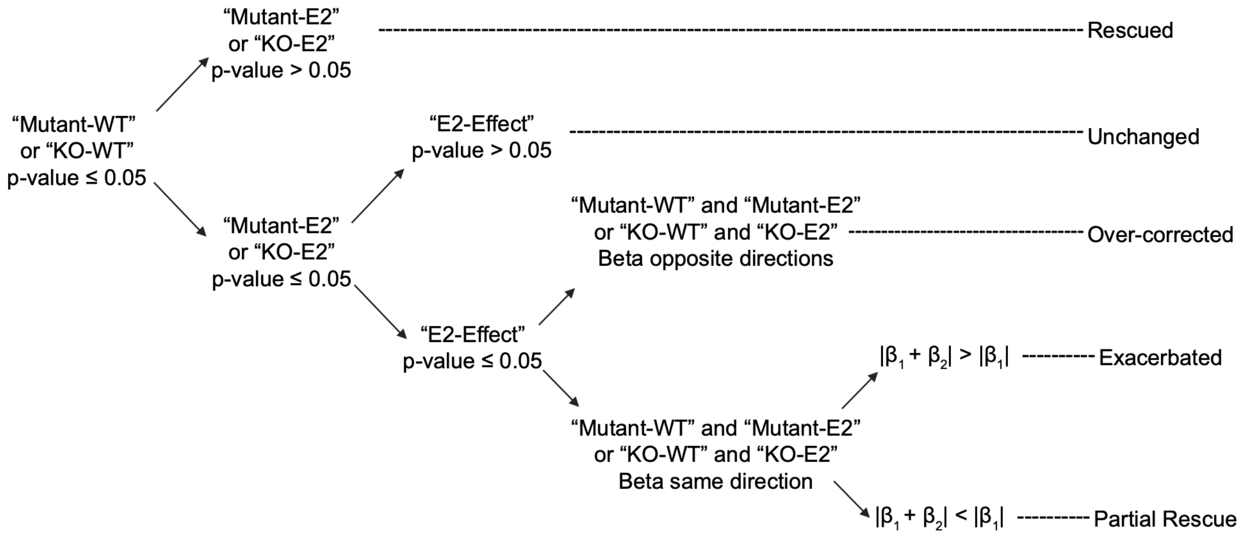


***SI Figure 16: Degree of rescue thresholds criteria.*** Decision tree thresholds dictate for any given parameter the degree of estradiol rescue.
